# Continuous manipulation of mental representations is compromised in cerebellar degeneration

**DOI:** 10.1101/2020.04.08.032409

**Authors:** Samuel D. McDougle, Jonathan Tsay, Benjamin Pitt, Maedbh King, William Saban, Jordan A. Taylor, Richard B. Ivry

## Abstract

Here we test the hypothesis that the cerebellum aids in the dynamic transformation of mental representations. We report a series of neuropsychological experiments comparing the performance of individuals with cerebellar degeneration (CD) on cognitive tasks that either entail continuous, movement-like mental operations or more discrete mental operations. In visual cognition, individuals with CD exhibited an impaired rate of mental rotation, an operation hypothesized to require the continuous manipulation of a visual representation. In contrast, individuals with CD showed a normal processing rate when scanning items in visual working memory, an operation hypothesized to require the maintenance and retrieval of representations. In mathematical cognition, individuals with CD were impaired at single-digit addition, an operation hypothesized to require iterative manipulations along a mental number-line; this group was not impaired on arithmetic tasks requiring memory retrieval (e.g., single-digit multiplication). These results, obtained in tasks from two disparate domains, suggest one potential constraint on the contribution of the cerebellum to cognitive tasks. This constraint may parallel the cerebellum’s role in motor control, involving coordinated dynamic transformations in a mental workspace.

## INTRODUCTION

The functional domain of the cerebellum extends beyond sensorimotor control^1,2^. Anatomical studies have revealed broad patterns of connectivity between the cerebellum and most of the cerebral cortex in humans and non-human primates, including prominent reciprocal connections between the cerebellum and prefrontal cortex^3–5^. Work in rodent models has linked cerebellar activity to a variety of surprising non-motor functions, such as reward processing, decision-making, and social interaction^6–8^. Human functional neuroimaging studies have revealed consistent cerebellar activation patterns unrelated to overt movement^9–11^, and neuropsychological studies have identified a large set of non-motor tasks on which individuals with cerebellar pathology are impaired^2^.

The most common neuropsychological sequelae in individuals with cerebellar degeneration (CD) are impairments in cognitive control^12^, including visuospatial cognition^13^, working memory^14,15^, and abstract reasoning^16^. Echoing the loss of motor coordination observed in these patients (movement dysmetria), the phrase “dysmetria of thought” has been used to summarize heterogeneous cognitive symptoms associated with CD^17^. This phrase reflects the idea that core mental functions, such as perception and memory, are mostly spared in cerebellar pathology, but the ability to manipulate mental representations in a coordinated manner is compromised.

A number of computational hypotheses have been put forward based on the idea that cerebellar contributions to motor control may generalize to the cognitive domain ^18,19^. In the motor domain, computational accounts of cerebellar function have emphasized the importance of this structure in anticipating future states^20^. For example, in coordinating action, the cerebellum is hypothesized to generate predictions of the sensory consequences of a movement^21^. This prediction is compared with observed sensory feedback, with the difference used as an error signal to calibrate the sensorimotor system^22^.

Extending this idea to cognition has motivated hypotheses about a more general cerebellar role in prediction, and in particular, how the cerebellum may be required for mental simulation to predict future cognitive states given current inputs^19^. However, it is important to keep in mind that prediction is likely a general feature of brain function^23^; as such, a core challenge is to specify constraints associated with how a specific brain region or network contributes to prediction. In terms of motor control, one important constraint arises from the fact that movement entails the continuous transformation of the body: A movement goal may be couched in terms of a desired end state, with the motor system optimized to transform the body’s initial state to the desired end state in a smooth manner.

We ask here if this “continuity” principle provides a clue to understanding how the cerebellum contributes to cognition. Specifically, does the cerebellum assist in the continuous, coordinated transformation of mental representations? To test this idea, we draw on the cognitive psychology literature that places mental operations on a spectrum between those that emphasize continuous versus discrete transformations^24^. We hypothesize that the cerebellum might be especially important in tasks involving continuous, movement-like transformations of internal representations.

We tested the continuity hypothesis in two disparate non-motor domains, visual cognition and arithmetic. For visual cognition, an extensive body of behavioral and physiological research provides compelling evidence that operations involved in visual imagery and mental rotation entail continuous representational transformations^25^. For example, in mental rotation, the manipulation of a visual representation to facilitate object recognition entails movement through intermediate representational states^25,26^. This continuity constraint is strengthened by neurophysiological evidence, where movement planning elicits the continuous transformation of a population vector in motor cortex^27,28^. We hypothesized that individuals with CD would exhibit a disrupted (slowed) rate of mental rotation, which would be consistent with neuroimaging studies showing prominent cerebellar activation during mental rotation tasks^10,29^.

As a “non-continuous” control, we opted to use two visual memory search tasks in which participants compare a probe stimulus to a set of discrete items maintained in memory^28,30^. While recognizing that there are several differences between mental rotation and memory search, we chose the latter as a control because the rate-limiting factor in memory search does not require a continuous transformation of a single representation, but rather an iterative, retrieval operation on items held in working memory. We predicted that the CD group would not show a disrupted (slowed) rate of memory search.

To test the generality of the continuity hypothesis, we turned to a second domain, mental arithmetic, leveraging an analogous contrast between tasks that entail continuous transformations versus discrete retrieval. Multiple lines of behavioral and neuroimaging research posit the use of a continuous, spatialized representation that supports basic mathematical operations such as magnitude comparison and simple addition (i.e., a mental number line)^31–36^. In simple arithmetic tasks, reaction time increases in a linear manner with the magnitude of the operands^37^, and error patterns are consistent with having “momentum” while “moving” along the number line: People tend to overestimate solutions to addition problems and undersestimate solutions to subtraction problems^38,39^. Similarly patients with spatial neglect from right hemisphere lesions show systematic biases when mentally bisecting numerical intervals (e.g., stating 16 as midpoint between 12 and 18), similar to the rightward bias they exhibit in bisecting physical lines^40^.

These findings support the notion that simple addition entails a continuous mental transformation. As such, we predicted that individuals with CD would be impaired on a numeric verification task involving the addition of two single digit numbers and would show a steeper magnitude effect relative to controls. For our control tasks, we selected two numeric operations that have been demonstrated to rely on rote retrieval processes: Addition problems involving the same number (e.g., so-called “identity” problems such as 3 + 3) and single-digit multiplication (e.g., 5 * 6). For the former, reaction times are largely independent of the magnitude of the operand^41,42^. For simple multiplication, RT increases with magnitude, but the increase is attributed to a look-up operation mediated by problem frequency^43,44^. To the extent that identity addition problems and multiplication problems are memorized, solving them should not require a continuous transformation (e.g., “moving” four units along a spatialized number line to verify whether 3 + 4 = 7), but rather a discrete sampling of information from memory^45^. We thus predicted that performance on identity addition and multiplication would be similar in the CD and control groups.

## METHODS

### Participants

Adult participants diagnosed with spinocerebellar ataxia due to cerebellar degeneration (N = 48) and neurologically healthy controls (N = 42) participated in the study in exchange for monetary compensation ($20 per hour). All participants were screened for general cognitive deficits using the Montreal Cognitive Assessment. Inclusion required that the participant achieve a score above 20 on the 30-point scale. The protocol was approved by the institutional review boards at Princeton University and the University of California, Berkeley.

Inclusion in the CD group was based on either clinical diagnosis of cerebellar atrophy or genetic confirmation of a subtype of spinocerebellar atrophy (SCA) (see Table S1). 26 of the 48 individuals with ataxia had an identified subtype (SC1: 4; SCA2: 2; SCA3: 4; SCA5: 1; SCA6: 10; SCA8: 1; SCA15: 1; SCA28: 1; AOA2: 2); for the other individuals, genetic testing was inconclusive or absent. The 10 CD participants with confirmed SCA6 were all related. The CD participants were assessed at the time of testing with the Scale for Assessment and Rating of Ataxia (SARA^46^). Scores ranged from 2 (mild motor impairments) to 26 (severe motor impairments).

Control participants were recruited to provide a match to the clinical sample in terms of age, MoCA, and years of education (Table 1). The one exception was in Exp 1b; by emphasizing matches based on education in this study, we ended up with a CD sample that was significantly older than their control group. Given this difference, we included age as a covariate in the primary analyses of all the experiments (see below).

**Table 1:**
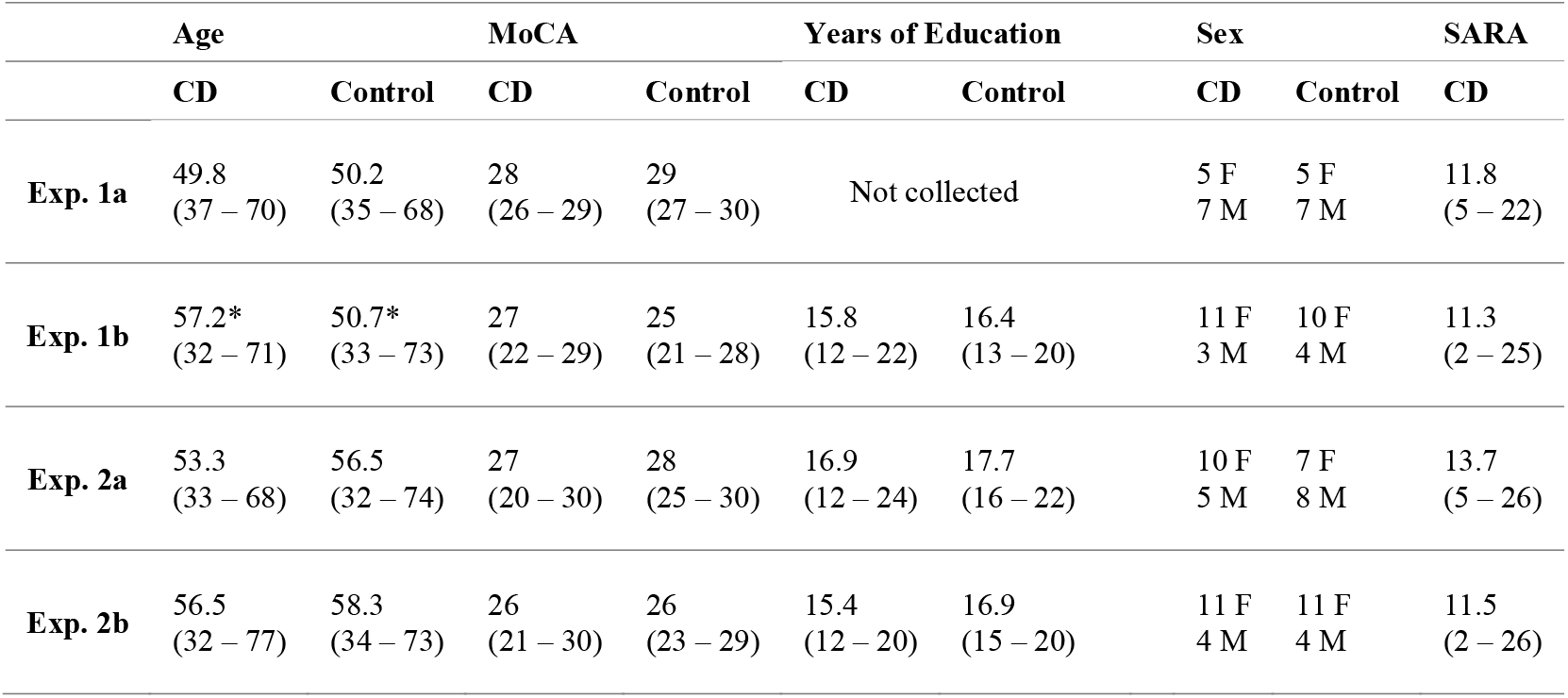
Demographic summary of CD and Control participants across all four experiments. Asterisks (*) denotes a significant difference (p < 0.05) between groups.

|  | Age |  | MoCA |  | Years of Education |  | Sex |  | SARA |
| --- | --- | --- | --- | --- | --- | --- | --- | --- | --- |
|  | CD | Control | CD | Control | CD | Control | CD | Control | CD |
| **Exp. 1a** | 49.8<br>(37 – 70) | 50.2<br>(35 – 68) | 28<br>(26 – 29) | 29<br>(27 – 30) | Not collected |  | 5 F<br>7 M | 5 F<br>7 M | 11.8<br>(5 – 22) |
| **Exp. 1b** | 57.2*<br>(32 – 71) | 50.7*<br>(33 – 73) | 27<br>(22 – 29) | 25<br>(21 – 28) | 15.8<br>(12 – 22) | 16.4<br>(13 – 20) | 11 F<br>3 M | 10 F<br>4 M | 11.3<br>(2 – 25) |
| **Exp. 2a** | 53.3<br>(33 – 68) | 56.5<br>(32 – 74) | 27<br>(20 – 30) | 28<br>(25 – 30) | 16.9<br>(12 – 24) | 17.7<br>(16 – 22) | 10 F<br>5 M | 7 F<br>8 M | 13.7<br>(5 – 26) |
| **Exp. 2b** | 56.5<br>(32 – 77) | 58.3<br>(34 – 73) | 26<br>(21 – 30) | 26<br>(23 – 29) | 15.4<br>(12 – 20) | 16.9<br>(15 – 20) | 11 F<br>4 M | 11 F<br>4 M | 11.5<br>(2 – 26) |

The test session last approximately 2 hours, including time for obtaining consent, medical history, performing the neuropsychological (all participants) and neurological (for CD only) evaluations, and conducting the experimental tasks. Breaks were provided between the different components of the session as well as partway through the experimental tasks. A test session included two different experimental tasks. A subset of the participants (CD = 8; Controls = 7) completed two of the tasks reported here Exps 1b and 2b) in a single session. For the other participants, the second task focused on sensorimotor learning.

### Apparatus and Procedure Overview

For all experiments, the stimuli were displayed and responses recorded on a laptop computer (MacBook Pro, Apple) using the psychophysics toolbox package^47^ for MATLAB (MathWorks). Participants were seated a comfortable distance from the screen (viewing distance ~40 cm). Responses were made with the index and ring fingers of the right hand on the computer keyboard.

### Experimental Tasks

#### Experiment 1

Experiments 1a and 1b were designed to evaluate the continuity hypothesis in the domain of visual cognition. Each involved two conditions, an experimental condition hypothesized to entail a continuous operation and a control condition, hypothesize to entail a non-continuous, or discrete operation. In both Exps 1a and 1b, the same mental rotation task was used for the continuous condition. It was paired with a visual working memory task in Exp 1a and a visuospatial working memory task in Exp 1b. The continuous and control conditions were tested in separate experimental blocks within a single session, with the order counterbalanced. Each task took approximately 25 min to complete, and participants were given a 10 min break between conditions.

#### Mental Rotation (Experiments 1a and 1b)

Following the basic design of Shephard and Metzler^26^, participants judged if a visual stimulus was normal (“R”) or mirror-reflected (“Я”; Fig 1). Eight capitalized sans-serif (Helvetica font) letter stimuli were used, consisting of normal and reflected versions of the letters F, G, J, and R^48^. Letter stimuli were white and presented on a black background. To minimize eye movements while maintaining stimulus legibility, the stimuli were modestly sized (~4 cm^2^), of high contrast, and were presented at a central location on the monitor.

**Figure 1:**
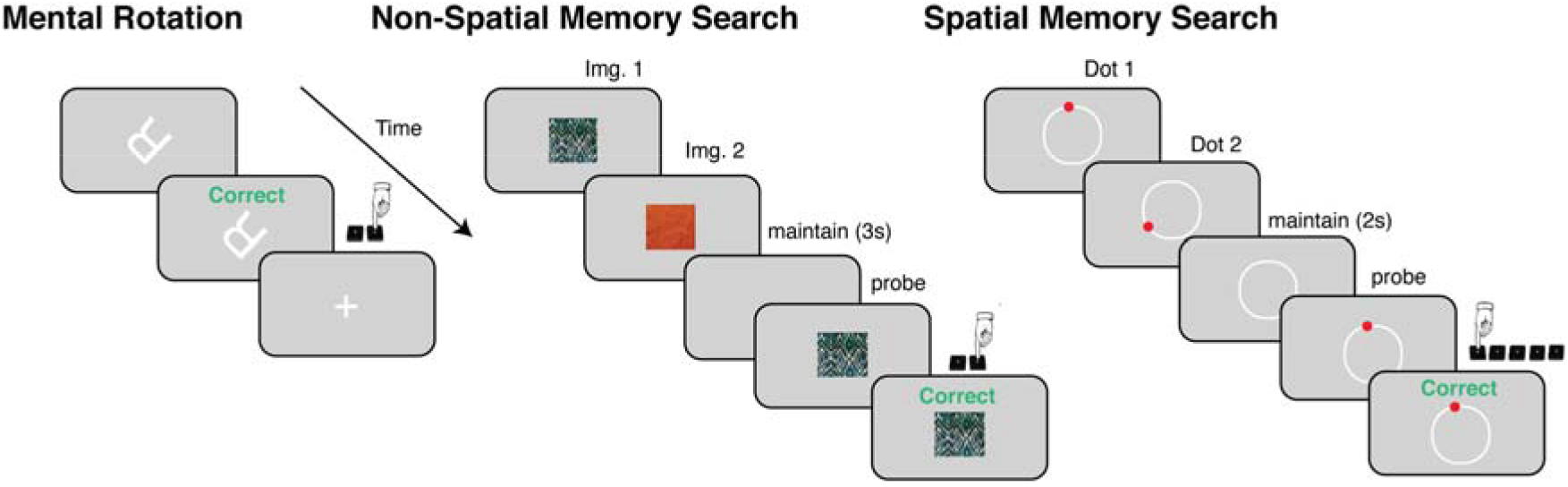
Visual cognition tasks employed in Experiments 1a and 1b. In the mental rotation experiment, participants judged if a letter stimulus was normal (e.g., “R”, right key press) or mirror-reflected (e.g., “Я”, left key press). On most trials, the stimulus was rotated relative to the upright orientation (depicted example involves 135° rotation). The same mental rotation task was used in Experiments 1a and 1b for the continuous condition. Two memory search tasks were used for the non-continuous, control conditions. In the visual memory search task (Exp 1a), a sequence of stimuli (1-5 images) was presented (1 s per image). After a maintenance period (3 s), a probe stimulus appeared, and the participant judged whether it was a member of the memory set (right key) or not (left key). Sequences varied in length from one to five items. In the visuospatial working memory task (Exp 1b), a sequence of circles (2-5 items) was presented at random locations on a ring (1 s per target). After a maintenance period (2 s), a probe stimulus was presented, and the participant indicated the ordinal position of the probe. Responses in all tasks were followed by feedback (1 s), and a 2 s inter-trial-interval (ITI).

Participants were instructed to press the right arrow key with their right ring finger when the stimulus (if viewed or imagined in an upright orientation) was in standard form and the left arrow key with their right index finger if the stimulus was mirror reflected. The stimulus was presented in the standard upright orientation (0°, baseline condition), or rotated, using one of 10 angles drawn from the following set: −135°, −105°, −75°, −45°, −15°, 15°, 45°, 75°, 105°, 135°. The stimulus remained visible until the response, or for a maximum of 5 s, whichever came first. After the response, feedback was shown above the letter for 1 s. On correct trials, the word “correct” was displayed in green font. On incorrect trials, the word “incorrect” was displayed in red font. Participants were instructed to respond quickly, while maintaining a high level of accuracy. If no response was made within 5 s, the message “too slow” was displayed in red font. Following a 1 s feedback interval, the display was replaced by a white fixation cross (0.9 cm^2^) that remained visible for a 2 s inter-trial interval.

Participants performed 18 trials at each rotation size and sign, intermixed with 36 no-rotation baseline trials, for a total of 216 trials. Stimuli were presented in a random order, with an equal number of normal and reflected presentations of each stimulus at each rotation sign and magnitude. Prior to the start of the experimental block, the participants performed five practice trials to ensure that they understood the task instructions and were comfortably positioned to respond on the keyboard.

### Visual Memory Search (Exp 1a)

As a control task in Exp 1a, we employed a variant of the “memory scanning” task introduced by Sternberg^30^. On each trial, participants viewed a brief sequence of visual stimuli and, after a maintenance period, judged whether a probe stimulus was a member of the previous sequence (match) or not (non-match). The stimuli consisted of 30 colorful fractal-like patterns, generated using the randomization function of ArtMatic Pro (www.artmatic.com). The images were cropped to be square-shaped and were matched in size to the mental rotation stimuli (4 cm^2^).

The memory set was presented sequentially with each fractal image in the set displayed at the center of the screen for 1 s (with no inter-stimulus interval). To vary the working memory load across trials, the number of stimuli in a set ranged from 1 to 5 items. After the sequence terminated, the screen was blanked for a maintenance period of 3 s. A probe stimulus was then presented. Participants were instructed to press the right arrow key with the right ring figure in the event of a match, and the left arrow with their right index finger in the event of a non-match. The probe remained visible until the response or until 5 s had elapsed. Feedback (“correct”, “incorrect”, “too slow”) was displayed above the probe stimulus for 1 s after the response was made. The display was then replaced by a white fixation cross for a 2 s inter-trial interval.

On 50% of the trials, the probe matched one of the items in the sequence, and on the other 50% of the trials, the probe did not match any of the items. Twenty trials at each set size were presented (10 match, 10 non-match) in a random order for a total of 100 trials. Participants completed five practice trials at the start of the experiment.

### Spatial Visual Memory Search (Exp 1b)

A spatial working memory task was employed for the control condition in Exp 1b, adopted from a task introduced by Georgopoulos and Pellizzer^28^. On each trial, a sequence of red circles (diameter 1.2 cm) was displayed on a white ring (radius 7 cm). A circle could be presented at any location from 0° - 345° (at multiples of 15°), with the constraint that no location be repeated in a given sequence. Each stimulus was presented for 800 ms, with no time gap between successive targets. The number of stimuli in the sequence ranged from 2 to 5 items. Following the offset of the last item in the sequence, the white ring remained on the screen for a maintenance period of 2 s, after which a probe stimulus was shown. The probe always appeared in one of the positions previously shown in the sequence. Participants were instructed to press the number on the keyboard corresponding to the ordinal position of the probe within the sequence (i.e., “1” key if location of first item, “2” key if location of second item, etc.). The probe remained visible until the response, and feedback was presented for 1 s following the response. During the 2 s inter-trial interval, the white ring remained visible.

Each set size (i.e., sequence length, 2 - 5) was presented 30 times in a randomized order, for a total of 120 trials. Within each set-size, probe positions were sampled uniformly between the first and the second-to-last position; for example, if the set size was 5, the probe location could match the location of the first, second, third, or fourth target in the sequence. Except for a set size of 2, we chose not to include trials probing the terminal position given the asymmetrically large RT benefit for this position observed in pilot testing. The task started with five practice trials to ensure that participants understood the instructions.

#### Experiment 2

Experiments 2a and 2b evaluated the continuity hypothesis in the domain of mathematical cognition. Exp 2a consisted of only addition problems where the continuous condition was composed of equations in which the two operands were non-identical, and the control condition was composed of equations in which the two operands were identical. These two types of equations were intermixed in a single block of trials that took approximately 25 min to complete. Both addition and multiplication problems were tested in Exp 2b, with the latter providing a second control condition. To minimize task-switching costs, the two types of mathematical operations were tested in separate blocks, each lasting approximately 25 min (plus the 10 min break), with the order counterbalanced across individuals.

### Addition Verification (Exp 2a and 2b)

On each trial, participants indicated if an addition equation, composed of two single digit operands and a sum, was true or false (Fig 4)^42^. The continuous condition consisted of trials in which the operands were non-identical (e.g., 4 + 7 = 11); the control condition consisted of trials in which the operands were identical (e.g., 6 + 6 = 12). Equations were white and presented at the center of the screen on a grey background. To minimize the necessity of making large saccades while maintaining stimulus legibility, the total length of the equation was modestly sized (4 cm). The equations were presented in a standard format (e.g., 3 + 7 = 11) with a double digit always used for the sum (e.g., “09” if the indicated sum was 9) to reduce the use of heuristics (e.g., recognizing that 3 + 1 could not be a two-digit sum).

Equations were drawn from a set of 36 single digit equations. Equations in Exp 2a were comprised of all unique combinations of operands between 3 and 8. Equations in Exp 2b were comprised of all unique combinations of operands 3, 4, 6, 7, 8, 9. Operands less than 3 were removed to limit the number of combinations where magnitude effects were modest^42^. We also excluded the operand 9 in Exp 2a to further limit the number of combinations; we replaced the 5 with 9 in Exp 2b given concerns that multiplying by 5 (see Exp 2b control condition below) would be easier than other two-digit multiplication problems. Each equation was presented eight times, consisting of four true responses and four false responses. True responses had one unique equation with the correct sum provided (e.g., 3 + 4 = 7). For the false equations, there were four distinct erroneous sums for each equation, with the presented answer different from the actual sum by either ± 1 (e.g., 3 + 4 = 8; 3 + 4 = 6) or ± 2 (e.g., 3 + 4 = 9; 3 + 4 = 5).

There were 288 addition trials in total, with the equations with identical and non-identical operands randomly interleaved. The trial sequence was subject to three constraints to minimize the effects of numerical and response priming on reaction time^49^: 1) If answered correctly, the same response would not occur more than three times in succession; 2) Consecutive problems could not share identical operands; 3) Consecutive problems could not share the same solution. Note that this third constraint limits repetition of equations with the actual true solution, not the displayed solution, which could either be a true or false sum. The entire block took approximately 25 minutes.

#### Multiplication Verification (Exp 2b)

Participants completed a second block of trials in Exp 2b in which they performed the verification task on multiplication equations. This condition was added to provide a second control condition given the assumption that participants use a look-up table in computing the product of two single-digit numbers^41^. The method for this condition was identical to that used in the addition block, including the use of the same set of operands (3, 4, 6, 7, 8, 9). Equations with erroneous products also had four variants, here created by adding ± 1 to either the first operand (e.g., 8 × 3 = 27 or 8 × 3 = 21) or ± 1 from the second operand (e.g., 8 × 3 = 16 or 8 × 3 = 32). There was a total of 288 trials consisting of equations with identical and non-identical operands, which were analyzed separately as in addition. The order of the addition and multiplication tasks in Exp 2b was counterbalanced across individuals.

### Data Analyses

Trials associated with extreme outlier RTs (> ± 3.5 SD from the participant’s mean) were removed prior to the analysis of the RT data. Less than 2% of trials were removed for all tasks. Parametric assumptions were tested using the Shapiro-Wilk test for normality and Levene’s test for homogeneity of variance. When parametric assumptions were met (which was the case for the accuracy data), statistical tests were performed on the mean values; when these assumptions were violated (which was the case for RT data for both groups in all experiments), non-parametric permutation tests were employed. All statistical tests were conducted in R (GNU).

### Visual Cognition Tasks

The three visual cognition tasks were selected because each has been shown to produce a parametric function relating the main independent variable, absolute rotation magnitude in mental rotation or set size in the two working memory search tasks, to RT. Given this, we computed the slope of the RT functions using each participant’s raw RTs in a general linear regression model assuming gamma distributed residuals. This regression analysis was performed on correct trials only. We opted to use a model based on the gamma distribution given that this distribution has been shown to better approximate RT distributions compared to the normal distribution^50–52^.

Our hypothesis centered on the two-way interaction between group and task on RT slopes. Specifically, we predicted a larger slope for the CD group compared to the Controls on mental rotation, but no group difference on memory search. To directly compare group differences in RT slopes across tasks with different independent variables (degrees for mental rotation; set size for memory search), we first computed slope differences between groups for each task to obtain two effect sizes. We then took the difference between these two effects sizes and computed a p-value by comparing the actual difference in effect size against a permutation distribution obtained by shuffling group labels 1000 times. A p-value less than 0.05 would indicate a disproportionate group difference in RT slopes for mental rotation versus memory search. Two-tailed permutation tests were used to compare group differences in RT slopes, with alpha set at 0.05 (non-parametric tests were used due to failure to meet parametric assumptions).

As a secondary analysis we compared differences in the RT slopes for each task separately. To this end, we performed an ANCOVA, with age and baseline median-centered RT included as covariates. The latter was motivated by the finding that the CD group was slower in responding across all tasks and conditions (baseline RT operationalized as rotation = 0° in mental rotation, set size = 1 in visual memory search, set size = 2 in spatial visual memory search). To quantify the influence of symptom severity on RT slopes, we correlated RT slopes in the CD group against their SARA scores.

### Mathematical Cognition Tasks

With the exception of problems with identical operands, addition and multiplication verification tasks involving two single-digit operands have been shown to produce an increase in RT as a function of magnitude, with magnitude defined in a range of ways (max operand, min operand, first operand, second operand, or the solution)^42,53,54^. We opted to use max operand as the independent variable for both the addition and multiplication tasks in the present study, although the pattern of results was not changed when magnitude was operationalized with any of the four other definitions.

Unlike visual cognition, the same independent variable, max operand, applies across the mathematical cognition tasks. Therefore, performance on the continuous and control conditions could be directly compared using a linear mixed effect model. Raw RTs were entered into this linear mixed effect model (assuming normal distributed residuals, since models with gamma distributed residuals failed to converge), with fixed effects consisting of the max operand, group, problem type (Exp 2a: equations with identical vs non-identical operands; Exp 2b: equations with non-identical operands in addition vs non-identical operands in multiplication), age, and years of education, as well as a random effect consisting of Participant ID. Our primary hypothesis centered on the three-way interaction among the fixed effects: Specifically, we predicted that the RT effect as function of max operand would be larger for the CD group only on the addition problems involving non-identical operands. This interaction was evaluated using a standard Chi-squared (*χ*^2^) likelihood ratio test.

As in Exp 1a and 1b, we also performed a more conservative non-parametric permutation test to evaluate group differences in RT slopes, limited to non-identical RT functions. Here we again used an ANCOVA, allowing us to evaluate whether group differences in RT slopes were influenced by differences in age, baseline median-centered RT (operationalized as the intercept of the non-identical RT function, i.e., max operand = 0), and years of education. To examine the influence of symptom severity on this measure, we also correlated RT slopes in the CD group against their SARA scores.

### Data Availability

The data that support the findings of this study are available from the corresponding authors upon request.

## RESULTS

### Experiment 1a

In Experiment 1 we tested the prediction that individuals with cerebellar degeneration would be selectively impaired on a visual cognition task that required the continuous transformation of a mental representation. For the continuous task (Fig. 1), we employed a classic mental rotation task^26^. Participants (n=12 CD and n=12 Control) judged if a visual letter stimulus was normal (“R”) or mirror-reflected (“Я”), where the stimulus was rotated by a particular degree on each trial: [−135°, −105°, −75°, −45°, −15°, 0°, 15°, 45°, 75°, 105°, 135°]. For the non-continuous task, we employed a variant of a classic memory search task^30^ presumed to require a search (iterative or parallel) through a set of discrete visual representations held in working memory. Participants viewed a single or sequence of abstract, visual fractal stimuli (set size 1-5), and, after a brief maintenance period, were asked to judge whether a probe stimulus was a member of the set (match) or not (non-match).

### RT effects

We first assessed whether the results with both groups were in accord with the classic finding observed with these two tasks, namely that RTs increase with the experimentally titrated independent variable – rotation magnitude for mental rotation and set size for visual memory search. Replicating these classic results, regression slopes on the RT data from the correct trials from both tasks were positive (all *p*_perm_’s < 0.005) for the control and CD groups (Fig 2): Participants took more time to respond for larger rotations and larger set sizes.

**Figure 2:**
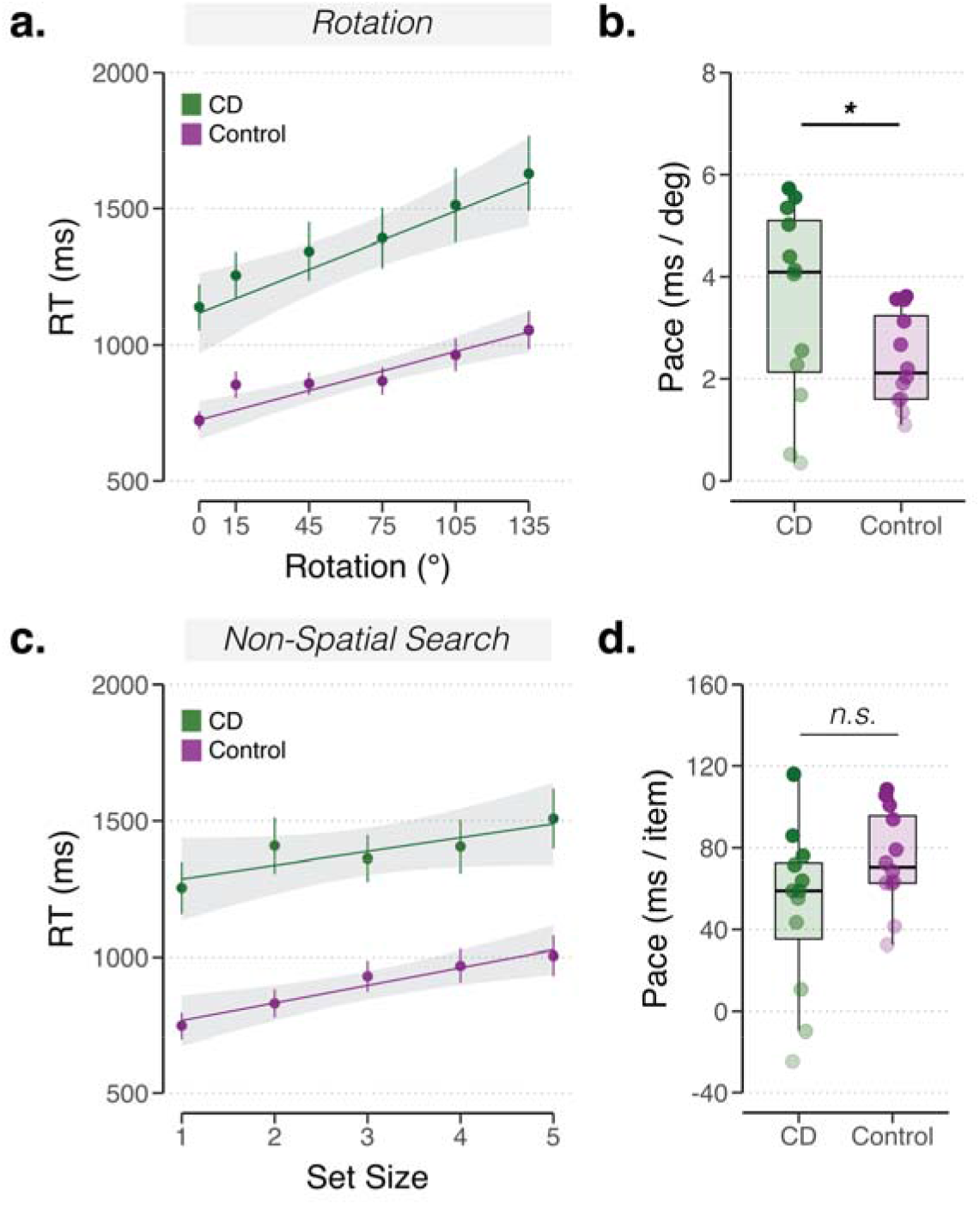
Reaction time analysis for Exp 1a. *Cerebellar degeneration is associated with a slower rate of mental rotation but does not impact the rate of search through visual working memory*. **(a)** Median RT as a function of stimulus orientation in the mental rotation task for the CD group (green) and the control group (purple). **(b)** Estimated rate of rotation from the regression analysis (slope of RT function). **(c)** Median RT as a function of set size in the visual memory search task. **(d)** Estimated search rate from the regression analysis. Mean regression lines are displayed in **(a)** and **(c)**. Shaded error bars denote 1 s.e.m. * *p* < 0.05.

We next compared behavior between the groups for each task separately. Collapsing over all correct trials on the mental rotation tasks, the CD group responded much slower than the Control Group (Mean ± SD: 1361 ± 600 ms vs. 869 ± 326 ms, *p*_perm_ = 0.006). The difference between groups may be due, at least in part, to the motor deficits associated with ataxia. However, it could also reflect differences in sensory encoding or a more generic impairment on performance associated with CD, an issue we return to in the Discussion.

Critically, our main interest is in the slope of the RT functions, a measure that indexes the rate of mental rotation (i.e., milliseconds per degree of rotation). An ANCOVA revealed a main effect of group, with the CD group showing a greater slope compared to the control group (*F*_(1, 20)_ = 4.5, *p* = 0.04). We interpret this increase in slope as indicative of a slower rate of mental rotation independent of any motor or generic impairment (given that these would be expected to impact RT independent of the angular shift of the stimulus). The slope effect persisted when controlling for the effects of age (*F*_(1, 20)_ = 0.8, *p* = 0.37), baseline RT (*F*_(1, 20)_ = 9.6, *p* < 0.001), and clinical ataxia ratings in the CD group (SARA scores; *R* = −0.1, *p* = 0.78) in the analyses. On average, the rate of rotation for the Controls was 2.2 ms/deg, whereas the rate for the CD group was 3.2 ms/deg. Using a more conservative non-parametric post-hoc assessment (two-sample permutation test, 1000 permutations), the CD mental rotation slopes remained significantly greater than Controls (*p*_perm_ = 0.01), corroborating the results from the ANCOVA.

At an individual level, 7 of the 12 CD participants had slower mental rotation rates than the slowest of the controls. Interestingly, two individuals in the CD group showed the fastest mental rotation rates (lowest slopes) overall (Fig 2b, two lightest dots in the CD group). These two participants also had the highest error rates on the task, responding correctly on fewer than 75% of trials (Fig S1a). Although speculative, it may be that the difficulty these individuals had in mental rotation led them to use an alternative strategy, making intuited responses based only on the presented orientation of the stimuli. Unsurprisingly, if these two participants were excluded from the slope analysis the resulting group difference remained significant (*p*_perm_ = 0.004).

Turning to the memory search task, we again observed a group difference in RT (CD: 1385 ± 560; Control: 889 ± 348, *p*_perm_ = 0.004), with the CD group responding slower overall. Critically, in contrast to the mental rotation results, the effect of group was not significant on the RT slope estimates (*F*_(1, 20)_ = 3.1, *p* = 0.09). Although this *p* value approaches significance, it is in the opposite direction, such that the average memory search rate for the CD group was faster than the average rate for the Controls (51 ms/item in CD vs 74 ms/item in Controls). The results of a two-sample permutation test failed to reveal a difference in slopes between groups (*p*_perm_ = 0.16). Moreover, we did not observe group differences in RT slopes when controlling for the effect of age (*F*_(1, 20)_ = 1.5, *p* = 0.24) and baseline RT (*F*_(1, 20)_ = 0.9, *p* = 0.30). The RT slopes of the CD group were again not correlated with SARA scores (*R* = −0.1, *p* = 0.83).

To directly compare RT slopes across the two tasks with different independent variables (rotation degrees vs set size), we used a permutation test on the group-level effect sizes. This comparison revealed a significant interaction between group and task, with the rate difference between groups being much larger for mental rotation compared to visual memory search (*d_Rotation_* − *d_Search_* = 1.45, *p_perm_* < 0.001). This dissociation suggests that the slower pace of rotation observed in the CD group did not reflect a global impairment in rate-based cognitive processing. Rather, the analyses point to a selective impairment in the ability of the CD group to perform mental rotation, consistent with the hypothesis that the integrity of the cerebellum is essential for efficiently coordinating the continuous transformation of an internal representation.

### Accuracy effects

Although our predictions focused on RT, we also examined accuracy on the two tasks (Fig S1a). We anticipated that overall accuracy would be lower for the CD group on both tasks, a common observation with this group (and most neurological groups) on speeded tasks^55^. Performance across both groups was higher on the mental rotation task (93.2%, SD = 7.2) than on the memory search task (85.7%, SD = 6.7). The control group had higher accuracy scores than the CD group in both the mental rotation s(*F*_(1, 20)_ = 5.7, *p* = 0.03) and memory search tasks (*F*_(1, 20)_ = 19.8, *p* < 0.001). Accuracy in the mental rotation task was affected by age (*F*_(1, 20)_ = 1.3, *p* = 0.27) and baseline RT (*F*_(1, 20)_ = 10.3, *p* = 0.004); however, accuracy in the memory search task was not affected by either variable (age: *F*_(1, 20)_ = 1.2, *p* = 0.28; baseline RT: *F*_(1, 20)_ = 0.04, *p* = 0.85). Task accuracy in the CD group was not significantly correlated with SARA scores (MR: *R* = −0.1, *p* = 0.53; MS: *R* = −0.56, *p* = 0.06).

Given the group difference in accuracy on both tasks, it is important to consider whether the dissociation observed between the tasks in the RT slope analyses might arise from a difference in the trade-off between speed and accuracy. If accuracy slopes between groups were similar, a speed-accuracy account of the RT slope analyses would be less likely. To examine this question, we performed another set of regressions using accuracy as the dependent variable (Figs S1b, c). Rotation magnitude and set size were associated with decreases in accuracy. However, the slopes were not different between groups (rotation: *p*_perm_ = 0.77; search: *p*_perm_ = 0.50; task X group interaction: *d_Rotation_* − *d_Search_* = −0.13, *p_perm_* = 0.59). Thus, the regression analyses provide no indication that the CD impairment in accuracy became more pronounced, relative to Controls, with larger rotations or increases in set size, arguing against a speed-accuracy tradeoff account of the RT results.

### Experiment 1b

The main factor in selecting the control task in Exp 1a was to have an independent variable – set size – that would produce a parametric increase in RT, providing a rate measure to index an iterative mental operation performed on a set of discrete representations. However, we recognize that there are many differences between mental rotation and visual memory search. While many of these differences are factors thought to affect the intercept of the RT function (e.g., the decision boundary for making the 2-choice speeded decision), one notable difference in the two tasks is that spatial representation is much more central in the mental rotation tasks. Given the association of the cerebellum in spatial processing, be it for motor control or spatial cognition^56^, we conducted a second visual cognition experiment to ask if the results in Exp 1a reflect a selective role for the cerebellum in spatial cognition rather than a selective role in facilitating continuous representational transformations.

To compare these two hypotheses, we paired the mental rotation task with a new control task designed to tax visuospatial working memory (Fig 1). Here, a sequence of circles was presented on a visual ring, and after a delay period a probe stimulus was displayed, where the position of the probe matched the position of one of the previously viewed circles. The participant indicated the ordinal position of the probe within the observed sequence by pressing one of five numbered keys. By varying the length of the sequence (2-5 locations), RT was expected to increase with set size, presumably reflecting a search through a set of discrete visual representations held in working memory. If the dissociation observed in Exp 1a is related to the spatial processing demands associated with mental rotation rather than the continuous nature of the required mental transformation, we would expect to observe an increase in slope in the CD group relative to the control participants on both the mental rotation task and the spatial working memory search task. In contrast, the continuity hypothesis would be supported if the CD group showed a selective slope effect on the mental rotation task. 14 participants with CD and 14 control participants were tested in Exp 1b, none of whom had participated in Exp 1a.

### RT effects

Rotation size and set size induced an increase in RT in both groups: RT slopes were positive for both groups in the mental rotation and working memory search tasks (Fig 3; all *p’s*_perm_ < 0.001). Like Exp 1a, RTs were significantly higher in the CD group compared to the Controls in both the mental rotation task (CD = 1365 ± 569 ms; Control = 1052 ± 527 ms, *p*_perm_ = 0.008) and in the memory search task (CD = 1566 ± 607 ms; Control = 1184 ± 542 ms, *p*_perm_ = 0.02).

**Figure 3:**
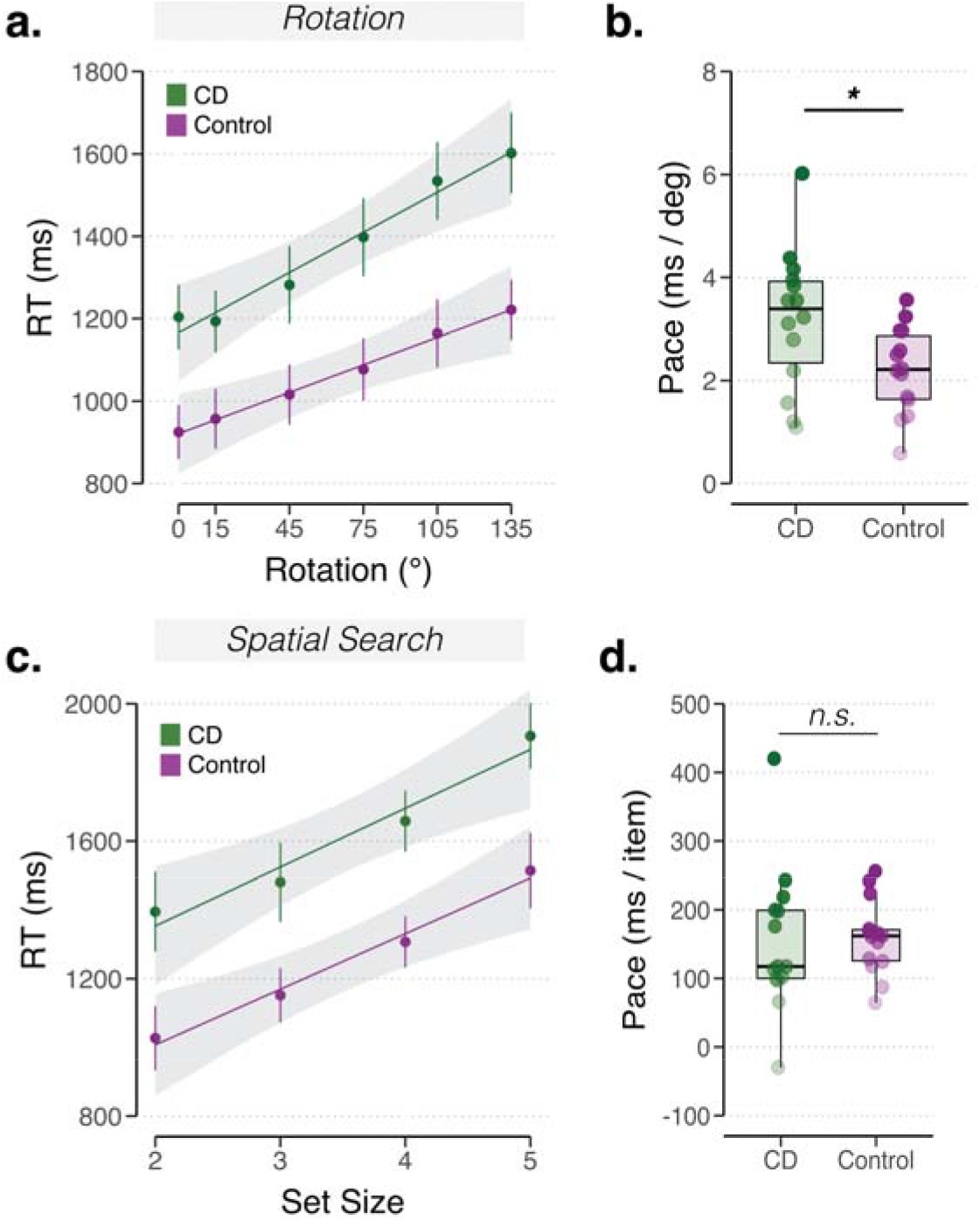
Reaction time analysis for Exp 1b. *Replication of selective impairment on the continuous transformation condition.* In mental rotation task **(a-b)**, CD group (green) showed slower mental rotation speeds relative to controls (purple). Mental rotation rates are plotted with the absolute rotation magnitude of the stimulus on the x-axis, and the median change in RTs for each rotation condition on the y-axis. In the search task **(c-d)**, the two groups showed comparable spatial memory search speeds. The length of the test sequence (set size) is plotted on the x-axis, and the median change in RTs for each set size on the y-axis. Mean regression lines are displayed in **(a)** and **(c)**. Shaded error bars denote 1 s.e.m. * *p* < 0.05.

**Figure 4:**
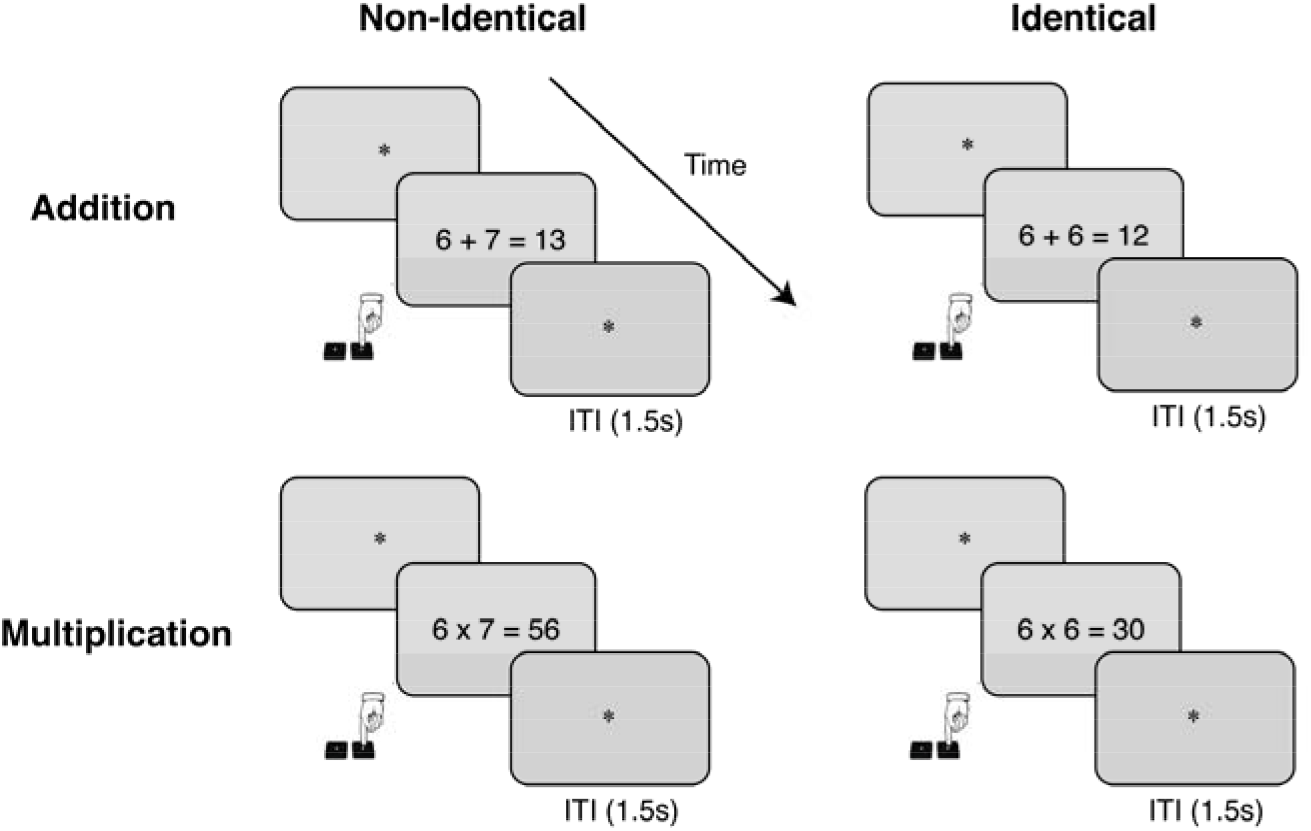
Mathematical cognition tasks employed in Experiments 2a and 2b. Participants made a speeded response to verify whether the equation was true or false. Addition problems (top row, tested in Exp 2a and 2b) and multiplication problems (bottom row, tested in only Exp 2b) involved either non-identical operands (left column) or identical operands (right column). Non-identical addition constitutes the continuous condition, whereas the remaining three conditions constitute non-continuous conditions.

Critically, we replicated the between-group slope difference on the mental rotation task: There was a main effect of group, with the CD group showing a higher slope compared to the control group (*F*_(1, 24)_ = 6.8, *p* = 0.03, Fig 3a, b). This difference was also observed in the direct two-sample permutation test (*p*_perm_ = 0.03). These group differences in mental rotation slopes persisted when controlling for age (*F*_(1, 24)_ = 0, *p* = 0.9) and baseline RTs (*F*_(1, 24)_ = 0.7, *p* = 0.4). In the CD group, RT slopes again did not correlate with SARA scores (*R* = 0.4, *p* = 0.16).

As a secondary analysis, we compared mental rotation RT slopes across the two experiments. The rotation rates were similar to those observed in Exp 1a with this new sample, with mean rates of 2.4 ms/deg for the Control group and 3.5 ms/deg for the CD group. A post-hoc, between-experiment comparison of the mental rotation slopes showed no difference in both CD groups (*t*_(24)_ = 0.09, *p* = 0.9; JZS Bayes Factor = 3.57 in favor of the null) and Control groups (*t*_(24)_ = −0.18, *p* = 0.9; JZS Bayes Factor = 3.53 in favor of the null), signaling a successful replication.

In contrast to the mental rotation task, there was no group effect of slope in the spatial memory search task, either when analyzed with the ANCOVA (*F*_(1, 24)_ = 0.1, *p* = 0.80, Fig 3c, d) or the direct two-sample permutation test (*p*_perm_ = 0.85). Thus, the CD group did not exhibit an increase in terms of the time required to search through a discrete set of spatial representations in working memory. Furthermore, we did not observe group differences in RT slopes when controlling for age (*F*_(1, 24)_ = 0.3, *p* = 0.58), baseline RTs (*F*_(1, 24)_ = 0.7, *p* = 0.58), and SARA scores in the CD group (*R* = 0.3, *p* = 0.25).

In a direct comparison of the rate measures between the two tasks, the CD group showed a significantly larger increase in slope on the mental rotation tasks in comparison to the visuospatial memory search task (task X group interaction: *d_Rotation_* − *d_Search_* = 1.45, *p_perm_* < 0.001). In summary, the RT data replicate the dissociation observed in Exp 1a: Degeneration of the cerebellum was selectively associated with a slower rate on a task that required a continuous transformation, but not on a task that involved an iterative search over a set of discrete representations. This dissociation is at odds with the hypothesis that the involvement of the cerebellum in visual cognition is tied to operations that require spatial processing.

### Accuracy effects

In terms of accuracy (Fig S2), performance was higher on the mental rotation task (94.2%, SD = 4.29) compared to the memory search task (77.8%, SD = 11.26). While this may reflect differences in task difficulty, it is important to note that in Exp 1b the number of response options differs for the two tasks, and thus the chance performance level (two options in mental rotation, chance level is 50%; five in memory search, chance level between 50% - 20% depending on set size). Unlike Exp 1a, accuracy scores were not significantly different between the CD and Control groups on the mental rotation task (*F*_(1, 24)_ = 0.3, *p* = 0.57). However, the CD group performed worse overall that the Control group on the memory search task (*F*_(1, 24)_ = 5.0, *p* = 0.04). Accuracy on both tasks was not affected by age (rotation: *F*_(1, 24)_ = 0.8, *p* = 0.4; search: *F*_(1, 24)_ = 0.03, *p* = 0.87), baseline RT (rotation: *F*_(1, 24)_ = 0.2 *p* = 0.66; search: *F*_(1, 24)_ = 0.5, *p* = 0.48), or SARA scores in the CD group (rotation: *R* = 0.3 *p* = 0.29; search: *R* < 0.1, *p* = 0.93).

As in Exp 1a, we asked if the slope differences might reflect a speed-accuracy tradeoff (Fig S2b, c). As expected, the functions relating rotation magnitude and set size to task accuracy were negative. However, the slope of these functions did not differ significantly between groups (rotation; *p*_perm_ = 0.74; search; *p*_perm_ = 0.97; task X group interaction: *d_Rotation_* − *d_Search_* = 0.15, *p_perm_* = 0.44), inconsistent with a speed-accuracy tradeoff account of the RT data. In sum, while the group difference in accuracy on the spatial search task might reflect a role of the cerebellum in spatial working memory, the results from the primary dependent variable (RT slope) and speed-accuracy analysis suggest that this contribution is not related to the rate-dependent operation of searching through a set of discrete spatial representations.

### Experiment 2: Assessing cerebellar contributions to mathematical cognition

The results of Exps 1a and 1b are consistent with the continuity hypothesis, with the selective impairment on the rate of mental rotation assumed to reflect the involvement of the cerebellum in transforming the orientation of a stimulus held in working memory. Importantly, there was no impairment in the CD group in processing rates for both the non-spatial and spatial memory retrieval tasks, arguing against a more generalized impairment in visual cognition. To further test the generality of the continuity hypothesis, we turned to a different domain in experiment two: mental arithmetic.

As noted in the *Introduction*, a large body of behavioral and neuroimaging research supports a contrast between algorithmic and memory retrieval processes involved in simple arithmetic^31,33,57^. For example, when participants solve addition problems with non-identical operands, RTs increase with the size of the operands (and their sum); in contrast, RTs remain flat when solving addition problems with identical operands^42,58^. Addition of non-identical operands has been posited to involve a continuous transformation of a spatialized mental representation (i.e., movement along a mental number line), whereas addition of identical operands is thought to involve rote memory retrieval from an overlearned look up table^33,36,45^.

Based on the continuity hypothesis, we predicted that individuals with CD would be slower in performing calculations involving a mental number line. Specifically, we predicted that they would show an elevated magnitude effect such that the RT difference between the CD and Control group would become larger with the magnitude of the operands, mirroring the effect of rotation magnitude in experiment 1. As a control task, we included arithmetic tasks associated with retrieval from a look-up table.

### Experiment 2a

Participants included 15 individuals with CD and 15 Control participants. We used a verification task in which the participants made a speeded response to indicate if addition equations composed of two single-digit operands and a sum were true or false (Fig 4). The operands were between 3 and 8 and included all pairwise combinations and orders. Unlike Experiment 1, the continuous and non- continuous conditions here involved the same task: The continuous condition was composed of trials in which the two operands were non-identical (e.g., 5 + 8), problems that are presumed to be solved with reference to a mental number line, while the non-continuous condition was composed of trials in which the two operands were identical (e.g., 6 + 6), problems presumed to be relatively automatized and solved via memory retrieval^42,53^. The participants completed a single block that included 230 non-identical trials and 46 identical trials, with equations being true in half of the trials and false in the other half.

### RT Effects

The independent variable (operationalized as the max operand, see *Methods*) was identical for the non-identical and identical conditions. As such, we used linear mixed effects modeling to analyze both within-participant and between-participant fixed effects.

Compared to the Control group, overall RTs were slower in the CD group for both non-identical (CD = 2381 ± 1339 ms; Control = 1262 ± 757 ms, *p_perm_* = 0.002) and identical equations (CD = 1667 ± 833 ms; Control = 1054 ± 543 ms, *p_perm_* = 0.004). The effect of problem type (i.e., identical versus non-identical) was significant, (*χ*^2^(1) = 11.1, *p* < 0.001), indicating that evaluating equations with non-identical operands required greater processing times than identical operands. There was no interaction between group and problem type (*χ*^2^(1) = 2.9, p = 0.09). Thus, the overall RT advantage for the Controls was similar in the two conditions.

Our focus is on the change in RT that occurs as a function of the max operand. There was a main effect of max operand, with RT increasing with the max operand (*χ*^2^(1) = 65.2, *p* < 0.001). There was an interaction between problem type and max operand (*χ*^2^(1) = 22.6, *p* < 0.001) (Fig 5a & c): Whereas RT increased with max operand for the non-identical equations for both CD (*p_perm_* < 0.001) and Controls (*p_perm_* = 0.02), RT was essentially flat for the identical equations for both groups (CD: *p_perm_* = 0.07; Controls: *p_perm_* = 0.95). This pattern — positive slopes as a function of max operand for non-identical equations, and no slope for identical equations, is consistent with previous studies^42,53^.

**Figure 5:**
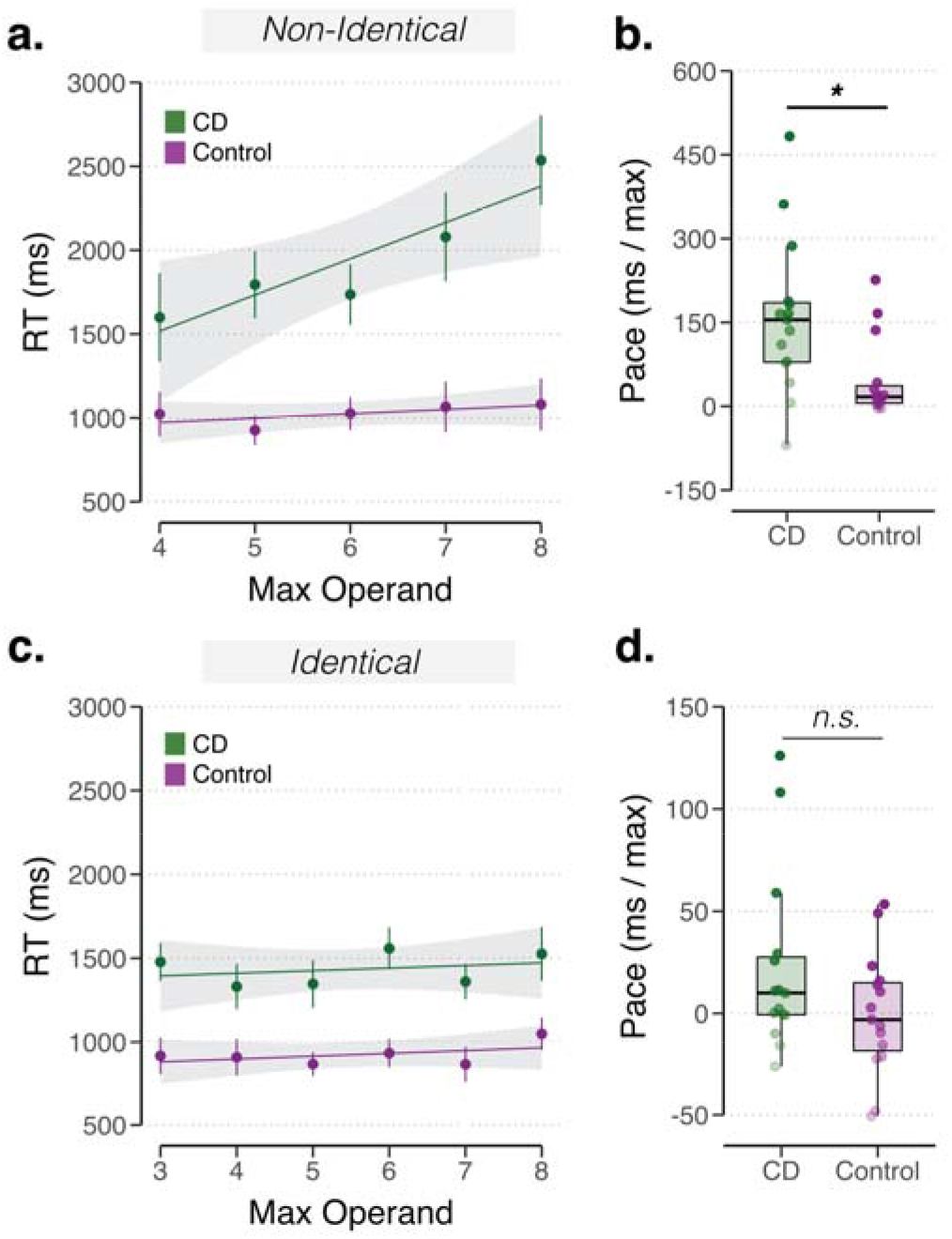
Reaction time analysis for Exp 2a. *Cerebellar degeneration associated with slower rate for addition problems putatively linked to continuous transformations along a number line, but not problems assumed to depend on memory retrieval*. Median RT is plotted as a function of the maximum operand for equations with non-identical **(a)** and identical operands **(c)**. Thin lines denote functions for each participant. The rate of adding non-identical **(c)** or identical operands **(d)**, is estimated by the slope of each individual’s RT function. Dots denote individuals. Mean regression lines are displayed in **(a)** and **(c)**. Shaded error bars denote 1 s.e.m. * *p* < 0.05.

Critically, this effect was modulated by group as indicated by the significant three-way interaction between the factors group, problem type, and max operand (*χ*^2^(1) = 18.1, *p* < 0.001). This three-way interaction in RT remained when we controlled for age (*χ*^2^(1) < 0.3, *p* = 0.60), years of education (*χ*^2^(1) = 2.6, *p* = 0.10), and SARA scores in the CD group (*R* = 0.12, *p* = 0.66). To analyze the three-way interaction, we estimated each individual’s RT slope using linear regression, similar to the procedure employed in Exp 1. Based on a direct permutation test, RT slopes when evaluating equations with different operands were found to be substantially larger in the CD group compared to Controls (Fig 5a; *p*_perm_ = 0.008). On average, individuals with CD required an additional ~150 ms per increment along the putative number line, whereas Controls only required an additional ~25 ms for each increment. In contrast, RT slopes for equations involving identical operands were similar between groups (Fig 5c; *p*_perm_ = 0.12; but see Exp 2b to address problematic nature of this comparison given that the slopes were not significantly different than zero for either group in this condition). Taken together, these results point to a selective impairment for the CD group on the addition task associated with a continuous operation rather than a general deficit in mental arithmetic.

### Accuracy Effects

As shown in Fig S3a, accuracy was high for both non-identical (94.77%, SD = 4.07) and identical equations (97.24%, SD = 3.40). The two groups performed equally well (*χ*^2^(1) = 0.2, *p* = 0.66), with all individuals performing above 80% on both problem types (Fig S3a-c).

There was a marginal effect of max operand on accuracy (*χ*^2^(1) = 3.2, *p* = 0.07), with accuracy tending to fall off with larger operands^42^. This fall-off was similar for both groups (*χ*^2^(1) = 0.2, *p* = 0.70), evidenced by the near-parallel lines seen in Figs S3b & 3c. Because the size of the operand had a similar effect on performance in both groups, a speed-accuracy trade-off cannot provide a parsimonious account for the RT results. There was no effect of age (*χ*^2^(1) = 1.65, *p* = 0.20), baseline RTs (*χ*^2^(1) = 1.0, *p* = 0.59), years of education (*χ*^2^(1) = 0.3, *p* = 0.59), nor SARA scores in the CD group (non-identical: *R* = −0.1, *p* = 0.62; identical: *R* = 0.34, *p* = 0.22) on accuracy.

### Experiment 2b

For the control condition in Experiment 2a, we selected equations involving identical numbers given prior evidence showing that these over-trained problems are generally solved via memory retrieval^42,53^. While this meets our objective to have a control task that involves a discrete operation, it is also suboptimal in one important way: RT did not increase as a function of the max operand when the equations involved identical operands. As such, this renders the rate measure difficult to interpret since the computation time remained invariant for all these equations (e.g., RT is the same for 2 + 2 and 6 + 6).

In Exp 2b we address this limitation by testing the participants on addition and multiplication problems, using the same set of operands for both tasks. The time required to perform and verify multiplication problems involving two single-digit numbers increases with the max operand, similar to that observed for non-identical addition problems^44^. However, the magnitude effect for multiplication has been associated with differences in the time required to access a look-up table, an effect related to the associative strength between operands and their solution, likely mediated by experienced problem^41,59^. Moreover, addition and multiplication have been linked to dissociable brain regions, with the former tapping into areas associated with spatial manipulation and the latter relying on areas more associated with verbal memory^33,45^. In using multiplication for the control task, we will be able to compare rate measures on the two tasks.

Consistent with the continuity hypothesis, we predicted that that the CD group would show an increase in slope for the addition problems, replicating Exp 2a, but not for the multiplication problems. An added feature of comparing addition and multiplication is that, *a priori*, we assumed that multiplying two single-digit numbers is more difficult than adding the same numbers. Thus, in this experiment, an impairment associated with CD is predicted for the easier task. 15 individuals with CD and 15 Controls were tested in Exp 2b (of these, 12 CD and 7 Controls also were tested in Exp 1b).

### RT Effects

We first consider the addition and multiplication equations with non-identical operands. Overall RTs in the CD group were slower than the Control group in both addition (CD = 1990 ± 897 ms; Control = 1305 ± 565 ms, *p_perm_* = 0.002) and multiplication equations (CD = 2346 ± 1269 ms; Control = 1505 ± 818 ms, *p_perm_* = 0.002), similar to the previous experiments. The effect of problem type on RT was not significant (*χ*^2^(1) = 1.8, *p* = 0.18). Critically, RTs increased robustly with the max operand for both groups (*χ*^2^(1) = 41.2, *p* < 0.001) in both problem types (problem type x max operand interaction not significant: *χ*^2^(1) = 0.3, *p* = 0.55). Thus, as predicted, both Control and CD groups showed positive RT slopes for addition (CD: *p_perm_* < 0.001; Control: *p_perm_* = 0.003) and multiplication (CD: *p_perm_* = 0.04; Control: *p_perm_* = 0.003).

There was a significant three-way interaction of group, problem type, and max operand (*χ*^2^(1) = 6.1, *p* = 0.01). This three-way interaction persisted when controlling for age (*χ*^2^(1) = 2.2, *p* = 0.14) and years of education (*χ*^2^(1) = 4.8, *p* = 0.03). Consistent with the continuity hypothesis, the increase in RT with increasing max operands was larger for the CD group compared to the Controls, but only for the addition condition (Fig 6a; *p*_perm_ = 0.011). Whereas the CD group required an additional ~65 ms for each integer increment in max operand on the addition problems, the comparable value for the Controls was only ~23 ms/integer. In contrast to addition, the RT slopes estimated from the multiplication condition were comparable for the two groups (Fig 6c; *p*_perm_ = 0.42). If expressed in the units used for addition, the mean rate values were 55 ms/integer and 48 ms/integer for the CD and Control groups, respectively. Importantly, the current results indicate that this putative memory retrieval operation was unaffected by cerebellar degeneration. We note that these null results held even when the three CD participants exhibiting negative slopes in their multiplication RT functions were removed (Fig 6d; *p*_perm_ = 0.69).

**Figure 6:**
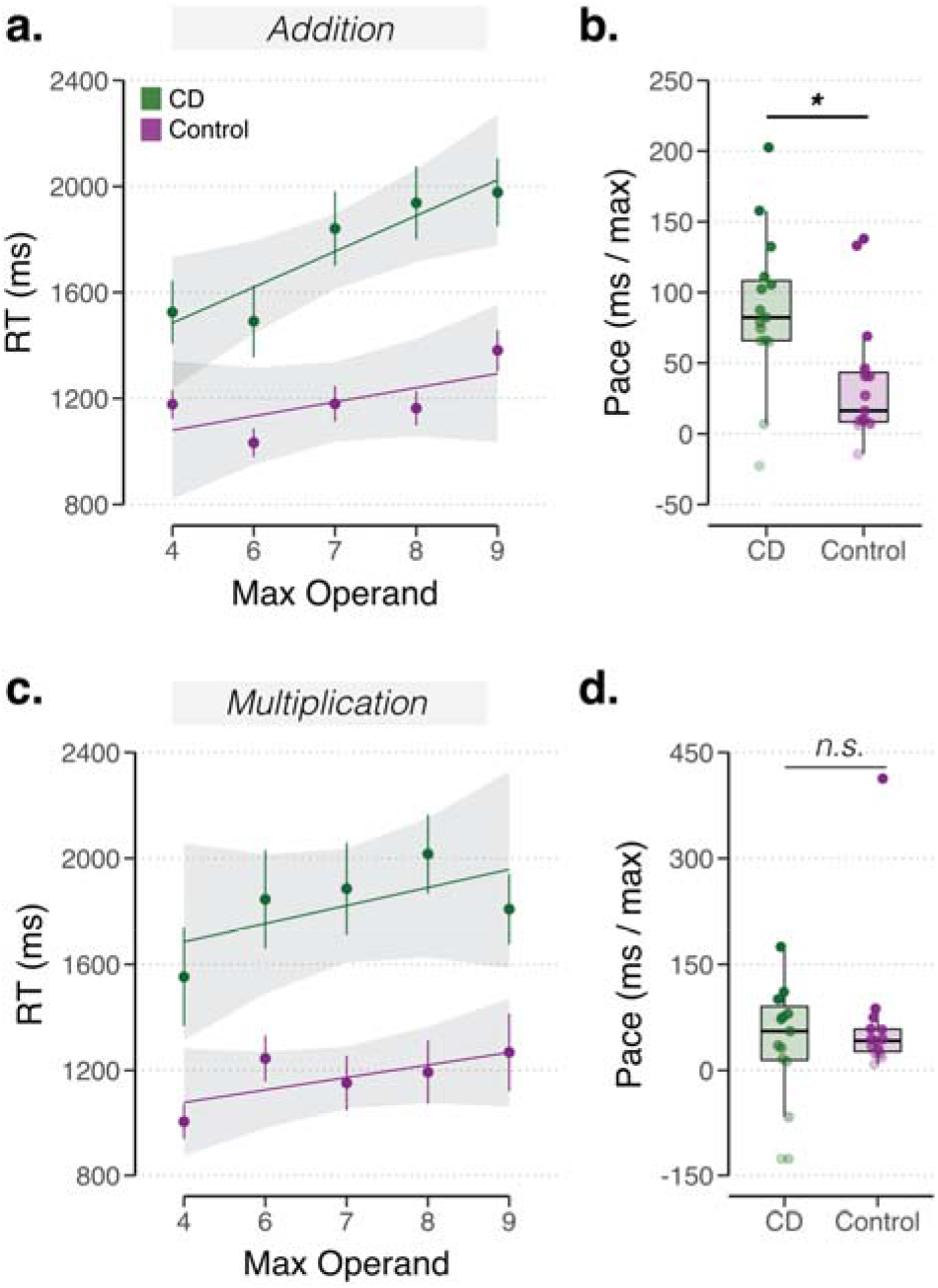
Reaction time analysis for Exp 2b. *Replication of selective impairment in CD group on arithmetic problems associated with continuous transformations along a number line.* Median RT is plotted as a function of the maximum operand for addition **(a)** and multiplication **(c)**, using equations with non-identical operands only. Thin lines denote functions for each participant. The rate for addition **(c)** or multiplication **(d)**, is estimated by the slope of each individual’s RT function. Dots denote individuals. Mean regression lines are displayed in **(a)** and **(c)**. Shaded error bars denote 1 s.e.m. * *p* < 0.05.

We performed a secondary, post-hoc analysis of the condition that was repeated in Exps 2a and 2b: addition of non-identical operations. This between-experiment comparison of the RT slopes for non-identical operands showed no difference in the Control groups (*t*_(28)_ = −0.6, *p* = 0.53; JZS Bayes Factor = 3.27 in favor of the null), signaling a successful replication. However, RT slopes in the CD group was significantly larger in Exp 2a compared to Exp 2b (*t*_(28)_ = 2.5, *p* = 0.02). While the reason for this difference is unclear, the larger slopes in CD groups compared to Controls in both experiments is consistent with the predictions of the continuity hypothesis.

### Accuracy Effects

As expected, there was a significant effect of problem type on accuracy (*χ*^2^(1) = 11.6, *p* < 0.001), with accuracy (Fig S4) lower on the multiplication problems (91.70%, SD = 5.52) compared to the addition problems (95.54%, SD = 0.03). Accuracy did not vary with age (*χ*^2^(1) = 2.8, *p* = 0.09) nor years of education (*χ*^2^(1) < 0.1, *p* = 0.93). Accuracy did co-vary with baseline reaction times, with lower accuracy associated with faster baseline reaction times (*χ*^2^(1) = 5.4, *p* = 0.02), an effect present in both Controls and CD groups. While there was no effect of the max operand on accuracy (*χ*^2^(1) = 2.19, *p* = 0.14), this variable interacted with problem type (*χ*^2^(1) = 30.4, *p* < 0.001). Accuracy declined at a faster rate for the multiplication problems compared to the addition problems (Figs S4b, 4c).

We again asked whether the RT effects might be mitigated by group differences in speed-accuracy tradeoffs. There was a significant three-way interaction in the accuracy data of max operand, group, and problem type (*χ*^2^(1) = 5.5, *p* = 0.02). Accuracy decreased faster as a function of max operand on multiplication problems for the CD group compared to the Controls, whereas the accuracy decrease with max operand was similar for the two groups on addition problems (multiplication: *p_perm_* = 0.03; addition: *p_perm_* = 0.87). This interaction could indicate that RT differences between groups for multiplication were masked by accuracy differences. We thus examined the relationship between the slope values for the accuracy and RT data: If participants were trading speed for accuracy, then a larger change in accuracy with max operand should be associated with a smaller change in RT with the max operand, and vice-versa. This correlation was neither observed for the Control group (multiplication: *R* = 0.03, *p* = 0.93; addition: *R* = 0.12, *p* = 0.7) nor the CD group (multiplication: *R* = 0.15, *p* = 0.56; addition: *R* = 0.4, *p* = 0.1). In sum, a speed accuracy account does not account for the null effects observed in the multiplication RT slopes.

### Comparison of problems with identical operands

While our main hypothesis centered on the comparison of addition and multiplication equations with non-identical operands, the addition and multiplication conditions included equations with identical operands. We directly extracted RT slopes using linear regression and compared these values between groups for each task. RT slopes were not significantly different between groups for addition with identical operands (*p_perm_* = 0.97) and multiplication with identical operands (*p_perm_* = 0.33). These results provide further evidence that cerebellar degeneration does not impact the speed of performing arithmetic operations assumed to be dependent on memory retrieval. Accuracy was very high in both tasks (identical addition: CD 96.94%, SD = 2.71%; Control 98.89%, SD = 1.55%; identical multiplication: CD 88.75%, SD = 9.12%; Control 94.44%, SD = 5.66%), and again, the observed functions did not suggest an elevated cost for the CD group for larger operands when the presented pair were identical (identical addition: *p_perm_* = 0.93; identical multiplication: *p_perm_* = 0.37).

### Comparison of subtypes of cerebellar degeneration

Extracerebellar pathology is often seen in some subtypes of SCA more than others; perhaps reflecting the extent of potential extracerebellar pathology, cognitive impairments on neuropsychological assessments are more common in some subtypes of SCA^60^. Although the sample size for different subtypes is small, the CD group did include 10 individuals with SCA-6 (all of whom were from the same family), a subtype in which the pathology is relatively limited to the cerebellum, and cognitive impairments tend to be minimal. As such, we performed follow-up exploratory analyses, comparing the individuals with SCA-6 to the other CD participants, focusing on Experiments 1b and 2b given the relatively balanced size of these two subgroups (Exp 1b: 5 SCA-6 vs 9 other subtypes; Exp 2b: 9 SCA-6 vs 6 other subtypes). No significant behavioral differences were observed: RT slopes did not differ between these two CD subgroups in either continuous operation condition (Exp 1b, mental rotation: *p_perm_* = 0.15; Exp 2b, non-identical addition: *p_perm_* = 0.75) or either discrete operation condition (Exp 1b spatial memory search: *p_perm_* = 0.25; Exp 2b non-identical multiplication: *p_perm_* = 0.47). While recognizing the limitations of our sample sizes for these analyses, the overall pattern suggests that the observed dissociation between the continuous and non-continuous tasks is likely associated with cerebellar degeneration.

## DISCUSSION

The cerebellum is a major anatomical structure of the central nervous system, strikingly conserved across the vertebrate subphylum. A large body of experimental and theoretical work has yielded detailed models of how this structure supports sensorimotor learning and motor control. In contrast, although the involvement of the cerebellum in cognition has been highlighted in many studies since the seminal conjecture of Leiner, Leiner, and Dow^1^, our understanding of its functional role has remained limited. The diverse patterns of task-related activity observed in neuroimaging studies of the human cerebellum might be taken to imply a heterogeneous role for the cerebellum in cognition^10,61^. Alternatively, the homogeneous anatomy and physiology of the cerebellum has inspired the idea that the cerebellum may invoke a common computation across diverse task domains^2,18^, or what has been called a “universal cerebellar transform” (UCT). By this view, the functional diversity inferred from the neuroimaging literature is seen as reflecting the diversity of inputs to the human cerebellum^62^, with a UCT being applied to these inputs to support a range of behaviors.

While recognizing that homogenous structure and physiology need not imply homogenous function, the UCT concept has been useful in generating testable computational hypotheses^19,22,63–65^. Here we build on this concept, seeking to identify constraints on the type of cognitive operations that rely on the cerebellum. Our theorizing draws on the distinction between continuous and discrete mental operations, a dimension that has proven useful in characterizing mental transformations^24^. The dynamics required to efficiently manipulate a body are inherently continuous, and the hallmark of cerebellar ataxia is a loss of fluidity in the movements required to transition from one state to another^20,22^. In a similar manner, we propose that the cerebellum might facilitate cognitive processing in tasks that involve a continuous representational transformation. This continuity hypothesis offers one potential constraint on the cerebellum and cognition: It specifies conditions under which the cerebellum would facilitate the efficient, coordinated operation of nonmotor mental operations and, as a corollary, conditions in which we would predict a reduced or negligible role of the cerebellum.

We evaluated the continuity hypothesis in two disparate domains. In the domain of visual cognition, we compared two classes of tasks: A mental rotation task known to be highly dependent on a continuous transformation^25,26,28^, and two variants of a visual memory search task presumed to require an iterative evaluation of the contents of a discrete set of representations in working memory. Reaction time slopes, taken as a proxy for the core mental operation required in these tasks, were elevated in the cerebellar degeneration group for mental rotation but neither for visual (Experiment 1a) nor visuospatial (Experiment 1b) working memory search. We note that this dissociation was observed in two independent samples of individuals with cerebellar degeneration (CD) and matched controls. Moreover, these results were robust to outlier and subgroup analyses (see *Results*).

To assess the generality of our hypothesis, we turned to a second domain, mathematical cognition. A distinction between continuous and discrete operations has also been articulated in this domain^31,36,38,42^. Drawing on this literature, we selected simple addition as a continuous operation given the evidence that this operation entails a translation across the internal spatialized representation of a number line^32,36,38,40^. For the discrete operation, we selected addition of identical single-digit integers or multiplication, operations hypothesized to rely on rote retrieval of instances from memory (e.g., 2 x 2). Consistent with our hypothesis, individuals with CD were selectively impaired on the simple addition problems, manifest as a larger increase in RT with operand magnitude compared to that observed in the Control group. In contrast, this slope difference was not observed on the two arithmetic tasks that depend on retrieval from a look-up table. As in Experiment 1, this dissociation was observed in two independent samples of participants and was robust to outlier and subgroup analyses.

We recognize that there are substantial differences in the continuous and discrete tasks we have employed in these experiments. This is especially true in Experiment 1, where we compare mental rotation with visual working memory tasks. Moreover, there are alternative ways to interpret the computational differences between the continuous and discrete tasks. Of note, the latter all involve some form of memory retrieval. We have opted to emphasize the continuous/discrete distinction. However, the current results would also be consistent with the idea that the cerebellum is not essential for the retrieval of information from memory but becomes essential when the retrieved information must be manipulated in working memory. Future research is required to pit this alternative hypothesis against the continuity hypothesis, or perhaps examine how the two hypotheses could be synthesized.

We note, though, that a retrieval-based account of the current results lacks parsimony on two fronts. First, retrieval in the current context requires lumping together two different types of retrieval: Retrieval of information just entered into working memory (Exp 1) and retrieval from long-term memory (Exp 2). Second, although the neuropsychological literature does indicate that individuals with SCA show minimal impairment on standard tests of memory^66^, these individuals tend to have difficulty on working memory tasks^16^. In contrast, the continuity hypothesis can parsimoniously explain the overall performance associated with SCA in two very different domains.

Another potential benefit of the continuity hypothesis is that it is offers a specific, falsifiable model of a cerebellar role in cognition. In this sense, we see it as a departure from previous conjectures that are more descriptive. For instance, while the concept of “dysmetria of thought” is an apt summary description of many previous clinical reports concerning cognitive deficits in cerebellar pathology^16^, it is unclear how this hypothesis can be operationalized. In contrast, the continuity hypothesis can be readily tested and falsified in other domains of cognition. For example, some models of visual attention posit a covert “spotlight” that traverses the visual scene^67,68^. Speculatively, we would predict that individuals with CD could be impaired at smoothly guiding a putative attentional spotlight through space, just as they are with guiding their eyes or limbs.

Our hypothesis also departs from previous theories that are more general in nature. While “prediction” has served as an important concept for understanding the role of the cerebellum in motor control^22^, it remains unclear how this concept should be applied to more cognitive domain, and more specifically, how the cerebellum’s predictive capacity might differ from predictive operations performed by the rest of the brain^23^. In motor control, this predictive capacity is thought to support the transformations needed to guide a limb in a continuous manner from position A to position B. Mental rotation could be described as a similar kind of continuous state transition problem, e.g., how do I, in my mind, rotate this object from orientation A to orientation B? Similarly, traversal of a mental number line would require an analogous state transition from one point in mental space to another (e.g., from left to right). On this view, our hypothesis shifts the focus away from a singular emphasis on prediction *per se*, and more towards consideration of the type of transformation required for a prediction-based computation.

The continuity hypothesis, however, cannot account for all of the behavioral results. First, the CD group was consistently slower to respond in all tasks and conditions. We assume some of the increase in RT is related to the global deficits associated with neurological disorder such as SCA (e.g., motor requirements for responding, even if these are simple movements; general resource allocation issues, etc.). However, baseline RT differences between the CD and control groups were around ~320 ms across the two mental rotation tasks and ~470 ms across the two working memory search tasks, values considerably larger than those observed in tasks involving relatively simple perceptual discriminations^69^. Second, although not dependent on set size, the CD group had worse accuracy than the control group on both of the memory search tasks (as well as on the mental rotation task). Taken together, these results again suggest that, in addition to a specific impairment in performing the continuous mental operations (i.e., the slope effects), the CD participants may also have more generic impairments that influenced their performance.

We also observed group differences in accuracy in the multiplication task (Experiment 2b), a deficit that became more pronounced as the operand increased. While subsequent analyses ruled out a speed accuracy account of the RT data, this accuracy effect is at odds with the continuity hypothesis, given the assumption that multiplication relies on memory retrieval^44^. We do not have a ready account of the accuracy effect in this condition. We note, though that, in designing these experiments, we deliberately avoided the use of accuracy as a primary dependent variable because this measure is likely to be sensitive to many cognitive processes: Poor accuracy can stem from attentional lapses, encoding errors, response biases, fatigue, etc., and all of these would be expected to compound with problem difficulty^55^. For these reasons, we selected tasks that allowed us to focus on parametric changes in RT (limited to correct trials), as this type of dependent variable provides a proxy of more specific cognitive operations (e.g., the rate of mental rotation, serial or parallel memory search, movement along a mental number line, etc.). We note that in two of our experiments (Experiments 1a and 2b), the tasks in which we did not see a CD deficit in RT slopes were apparently the more difficult of the two tasks, arguing against a generic difficulty confound.

The continuity constraint also offers a new lens to consider the results from previous studies examining the role of the in the cerebellum in visual cognition. The impairment we observed on mental rotation converges with prior work showing that deficits in visuospatial reasoning in individuals with cerebellar pathology (lesion or atrophy) are most pronounced on tasks requiring some sort of mental transformation^70^. More interesting is to consider the null results we report on the two visual working memory tasks in Experiment 1. At first blush, these findings appear at odds with the rather substantial neuroimaging literature pointing to the involvement of the cerebellum in working memory. Cerebellar activation is consistently observed in studies of spatial (and verbal) working memory^10,29,71,72^. However, the picture is less clear on the neuropsychological front, with some studies reporting impairments in individuals with cerebellar pathology on working memory tasks^15,73^ whereas others report null effects on these tasks^12,74^. Notably, the designs in these studies have tended to focus on main effects; for example, is activation higher on the working memory task relative to some baseline control, or do patients have reduced working memory span? The analyses typically do not examine rate-dependent measures, our chosen probe of individuals’ faculty for manipulating contents in working memory.

Considered more broadly, the continuity hypothesis can also be used to re-examine past work in other task domains. To give one example from our own work on temporal cognition, we have shown that individuals with CD are selectively impaired in using an interval-based representation for the production of periodic movement^75^ or temporal orienting^69^. In such tasks, performance requires reference to a continuously updated internal representation of time. In contrast, these individuals were unimpaired when temporal control can be achieved with a constant control parameter (e.g., angular velocity in tapping, entrainment to an exogenous referent signal in temporal orienting).

In summary, the continuity hypothesis provides a novel perspective on the contribution of the cerebellum to cognition, one that seeks to offer computational specificity to how the cerebellum facilitates mental coordination and prediction. By postulating constraints on cerebellar computation, we hope to advance our understanding of this important subcortical structure. Future studies involving a broader range of cognitive tasks will be essential to evaluate the utility of the continuity hypothesis as a candidate universal cerebellar transform.

## ACKNOWLEDGEMENTS

We thank the CognAc lab (UC Berkeley) and ACT lab (Yale) for helpful and thoughtful discussions.

## SUPPLEMENTAL MATERIAL

**Table S1:** *Demographic and clinical assessment of the CD participants.* MoCA scores are out of a total of 30 points. SARA scores are out of a total of 40 points (most severe ataxia). SCA = spinocerebellar ataxia; SAOA = spontaneous adult-onset ataxia.

| ID | Hand | Age | Sex | MoCA | SARA | Etiology | Years of Education | Exps Completed |
| --- | --- | --- | --- | --- | --- | --- | --- | --- |
| 1 | Right | 38 | F | 28 | 7.5 | SCA-1 |  | 1a |
| 2 | Right | 39 | M | 27 | 9.5 | SCA-1 |  | 1a |
| 3 | Right | 62 | M | 28 | 17.5 | SCA-1 |  | 1a |
| 4 | Right | 48 | F | 29 | 4.5 | SCA-1 | 24 | 2a |
| 5 | Right | 64 | F | 23 | 22.5 | SCA-2 | 16 | 2a |
| 6 | Right | 48 | F | 29 | 12.5 | SCA-2 | 16 | 2a |
| 7 | Right | 62 | F | 28 | 14.5 | SCA-3 | 16 | 1b, 2b |
| 8 | Right | 68 | M | 24 | 10 | SCA-3 | 18 | 2a |
| 9 | Right | 33 | F | 20 | 13.5 | SCA-3 | 12 | 2a |
| 10 | Right | 47 | F | 27 | 13.5 | SCA-3 | 16 | 2a |
| 11 | Right | 57 | F | 29 | 8.5 | SCA-5 |  | 1a |
| 12 | Right | 65 | F | 27 | 12.5 | SCA-6 | 22 | 1b |
| 13 | Right | 60 | F | 22 | 7.0 | SCA-6 | 12 | 1b, 2b |
| 14 | Right | 67 | F | 24 | 25.0 | SCA-6 | 12 | 1b, 2b |
| 15 | Right | 71 | F | 23 | 7.5 | SCA-6 | 16 | 1b, 2b |
| 16 | Right | 65 | F | 26 | 13 | SCA-6 | 16 | 1b, 2b |
| 17 | Right | 61 | M | 26 | 4.5 | SCA-6 | 16 | 1b, 2b |
| 18 | Right | 77 | F | 22 | 26 | SCA-6 | 12 | 2b |
| 19 | Right | 66 | F | 24 | 11.5 | SCA-6 | 17 | 2b |
| 20 | Right | 50 | F | 26 | 2 | SCA-6 | 12 | 2b |
| 21 | Right | 58 | F | 27 | 9 | SCA-6 | 12 | 2b |
| 22 | Right | 58 | M | 30 | 9 | SCA-8 | 20 | 2a |
| 23 | Right | 61 | M | 29 | 12.5 | SCA-15 |  | 1a |
| 24 | Right | 34 | M | 27 | 8 | SCA-28 | 20 | 2b |
| 25 | Right | 42 | F | 28 | 22.5 | AOA-2 | 18 | 2a |
| 26 | Right | 54 | M | 27 | 26 | AOA-2 | 14 | 2a |
| 27 | Right | 38 | F | 28 | 6 | SAOA |  | 1a |
| 28 | Right | 52 | M | 26 | 15.5 | SAOA |  | 1a |
| 29 | Right | 40 | F | 29 | 5 | SAOA |  | 1a |
| 30 | Right | 70 | F | 29 | 19.5 | SAOA |  | 1a |
| 31 | Left | 51 | M | 28 | 7.5 | SAOA |  | 1a |
| 32 | Right | 52 | M | 27 | 10.5 | SAOA |  | 1a |
| 33 | Right | 37 | M | 26 | 21.5 | SAOA |  | 1a |
| 34 | Right | 57 | F | 27 | 10.5 | SAOA | 18 | 1b |
| 35 | Right | 66 | F | 24 | 11.5 | SAOA | 17 | 1b |
| 36 | Right | 57 | F | 22 | 15.5 | SAOA | 12 | 1b |
| 37 | Right | 50 | M | 26 | 16 | SAOA | 20 | 1b |
| 38 | Right | 32 | M | 21 | 5 | SAOA | 16 | 1b, 2b |
| 39 | Right | 38 | F | 27 | 13.5 | SAOA | 16 | 1b, 2b |
| 40 | Right | 50 | F | 26 | 2 | SAOA | 12 | 1b |
| 41 | Right | 40 | F | 29 | 10 | SAOA | 14 | 2a |
| 42 | Right | 62 | M | 29 | 10.5 | SAOA | 16 | 2a |
| 43 | Right | 54 | M | 28 | 10 | SAOA | 16 | 2a |
| 44 | Right | 62 | F | 28 | 18 | SAOA | 14 | 2a |
| 45 | Right | 55 | F | 27 | 10.5 | SAOA | 18 | 2a |
| 46 | Right | 65 | F | 27 | 12.5 | SAOA | 22 | 2a |
| 47 | Right | 57 | F | 27 | 10.5 | SAOA | 18 | 2b |
| 48 | Right | 50 | M | 26 | 16 | SAOA | 20 | 2b |

**Figure S1:**
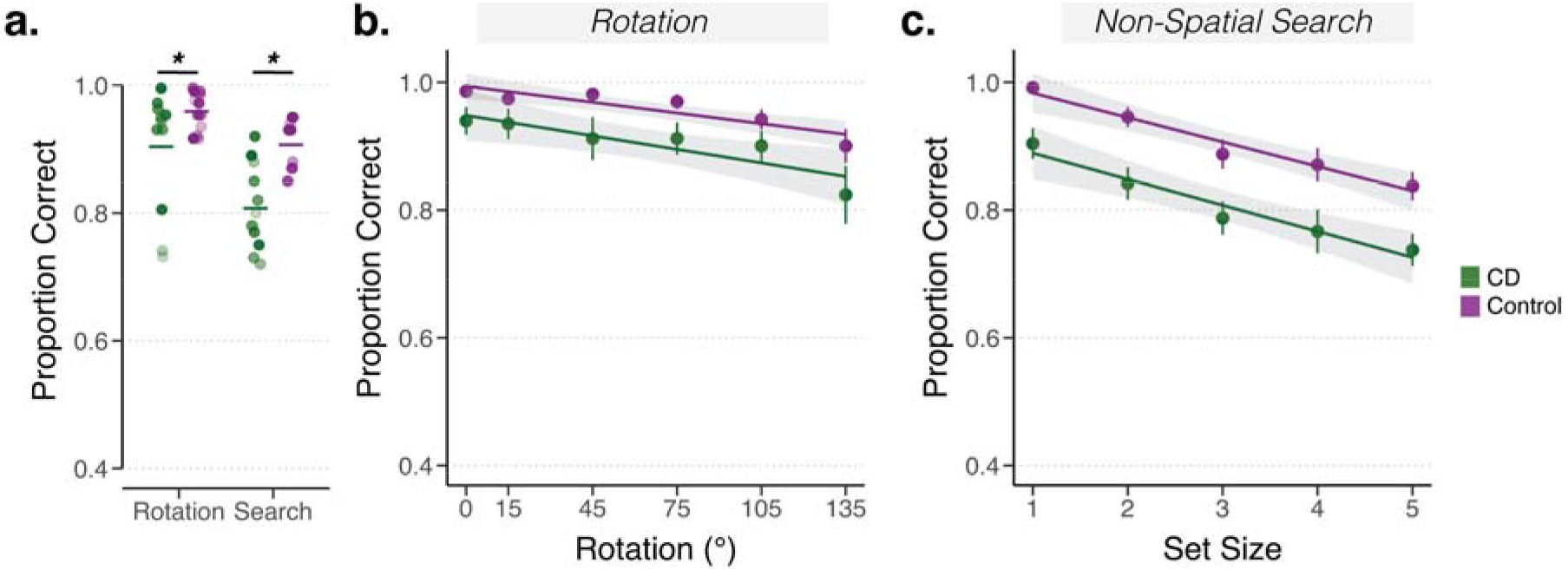
Accuracy analysis for Exp 1a. *Accuracy differences between groups are independent of rotation angle and set size in working memory.* **(a)** Overall accuracy rate is lower for the CD group compared to the Control group on both tasks (dots are shaded according to the within-group ranking of mental rotation and search rates, as seen in Figs 2b and 2d, with the darkest shade denoting the slowest rate and the lightest shade denoting the fastest rate; horizontal line indicates group means). Mean proportion correct as a function of rotation **(b)** and set size **(c)** for the rotation and memory search tasks, respectively. Mean regression lines are displayed in **(b)** and **(c)**. Shaded error bars denote 1 s.e.m. * *p* < 0.05.

**Figure S2:**
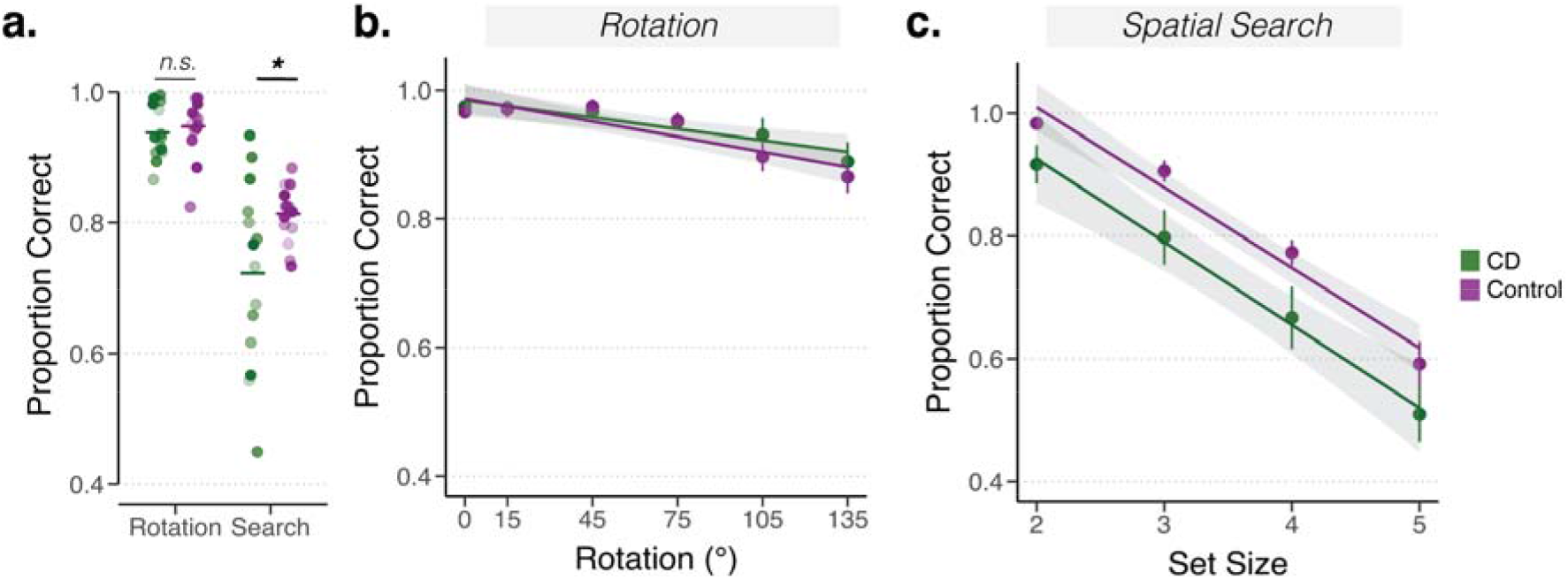
Accuracy analysis for Exp 1b. *Accuracy is a function of rotation angle and memory load, with group difference only observed on the search task.* **(a)** Overall accuracy rate is lower for CD group compared to Controls on the visuospatial working memory task (dots are shaded according to the within-group ranking of mental rotation and search paces, as seen in Figs 3b and 3d, with the darkest shade denoting the slowest rate and the lightest shade denoting the fastest rate; horizontal line indicates group means). Effects of independent task variables (rotation magnitude in panel **(b)** and set size in panel **(c)**) could not explain group differences. Error bars = 1 s.e.m. * *p* < 0.05.

**Figure S3:**
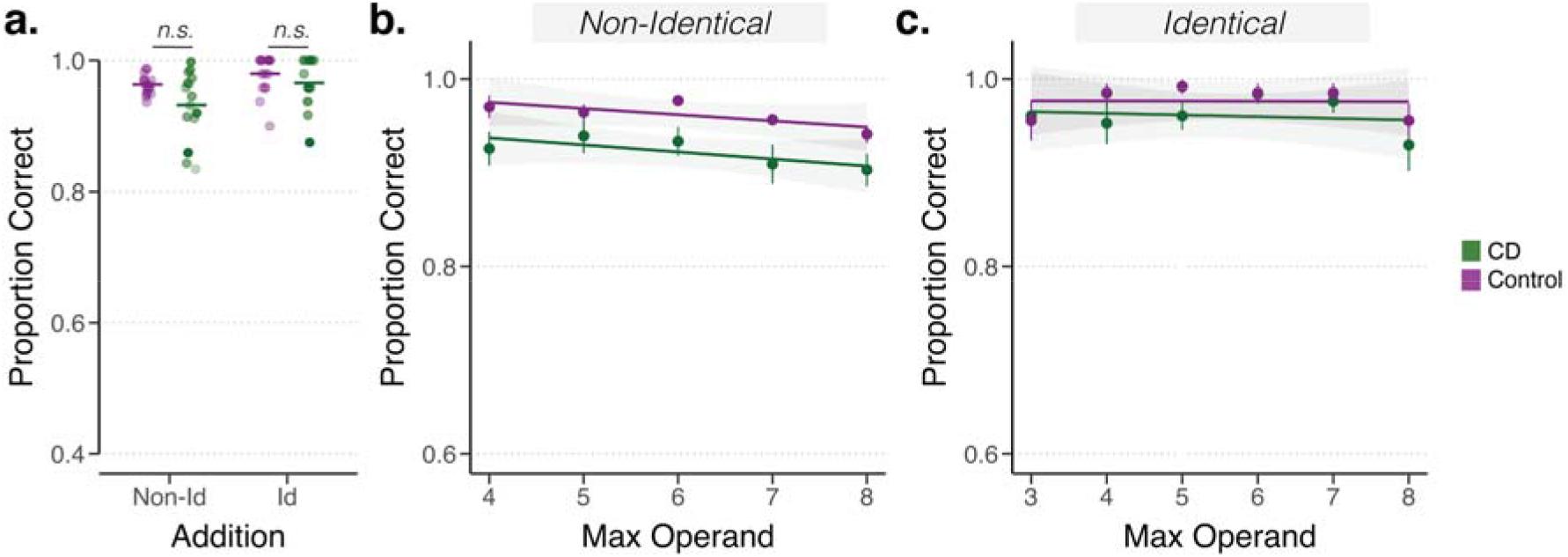
Accuracy analysis for Exp 2a. *Accuracy is comparable for the CD and Control groups in Exp 2a and does not provide evidence of a speed-accuracy tradeoff.* Overall performance **(a)** for each individual (dots) for the two types of addition problems (non-identical vs identical). Dots are shaded according to the within-group ranking of RT slopes, as seen in Figs 5b and 5d, with the darkest shade denoting the slowest rate and the lightest shade denoting the fastest rate; horizontal line indicates group means. Accuracy as a function of the maximum operand for equations with non-identical **(b)** and identical operands **(c)**. Mean regression lines are displayed in **(b)** and **(c)**. Error bars = 1 s.e.m.

**Figure S4:**
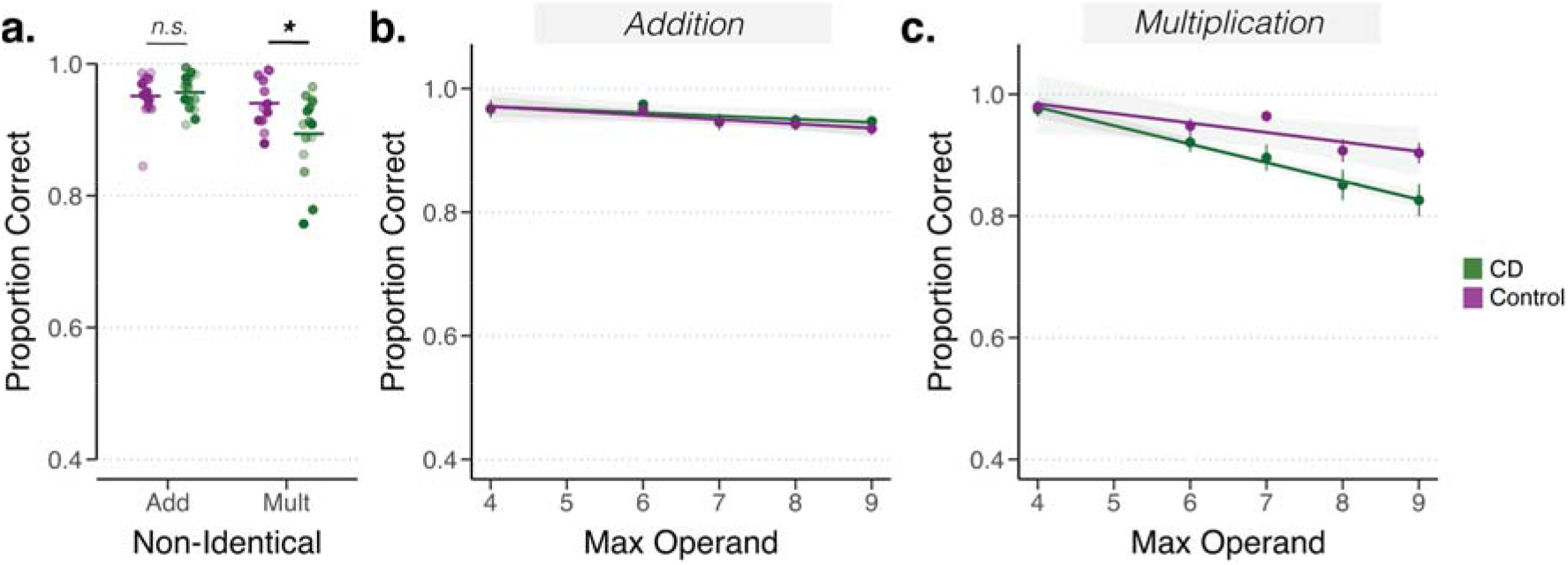
Accuracy analysis for Exp 2b. *Accuracy is comparable for the CD and Control groups and does not provide evidence of a speed-accuracy tradeoff.* Overall performance **(a)** for each individual (dots) for addition and multiplication problems consisting of non-identical operands. Dots are shaded according to the within-group ranking of RT slopes, as seen in Figs 6b and 6d, with the darkest shade denoting the slowest rate and the lightest shade denoting the fastest rate; horizontal line indicates group means. Accuracy as a function of the maximum operand for non-identical addition **(b)** and multiplication equations **(c)**. Mean regression lines are displayed in **(b)** and **(c)**. Error bars = 1 s.e.m.

